# Revealing the hidden sequence distribution of epitope-specific TCR repertoires and its influence on machine learning model performance

**DOI:** 10.1101/2024.10.21.619364

**Authors:** Sofie Gielis, Maria Chernigovskaya, Milena Pavlovic, Vincent Van Deuren, Romi Vandoren, Sebastiaan Valkiers, Kris Laukens, Victor Greiff, Pieter Meysman

## Abstract

Numerous efforts have been made to decipher the epitope-T cell receptor (TCR) recognition code. Both simple machine learning techniques and deep learning strategies have been used to train models to predict the binding of epitopes by TCR sequences. A good training data set rests at the basis of every accurate prediction model, yet little attention has been given to the composition of these data sets. In this paper, we studied the natural distribution of TCR sequences within epitope-specific TCR repertoires, i.e. a set of TCRs binding the same epitope, and its impact on the predictability of TCR-epitope interactions. We found that the observed diversity of these repertoires can result from a smaller set of core binding motifs constrained by TCR generation. Moreover, a clear relationship was found between the sequence distribution of the training data and performance metrics, emphasizing the importance of the used ground-truth data when using machine learning models in this domain. Taken together, these findings inform data set composition to help push epitope-TCR prediction models to the next level.

## Introduction

Our immune system contains various mechanisms to recognize and eliminate intruders. One mechanism is mediated by the molecular interaction between small peptides, i.e. epitopes, presented by the major histocompatibility complex (MHC) on the surface of nucleated cells, and T cells. These T cells are covered by epitope-specific receptors on their cell surface, the so-called T cell receptors or TCRs. The interaction between TCRs and MHC-presented epitopes activates the T cell resulting in a targeted immune response and the generation of memory T cells for faster activation in the future. TCR repertoires are collections of all sequenced TCRs per individual and thus directly capture information about current and previous immune states, including diseases, vaccinations and autoimmune reactions (1). Various attempts have been made to develop strategies to unravel epitope-TCR interactions from their amino acid sequences. Initially, these methods were focused on the identification of new TCRs recognizing a previously studied epitope (2–8). These prediction models show good performance for the majority of epitopes, provided that sufficient data is available. Due to the large diversity in epitopes, however, it is not feasible to collect and train individual models for each epitope in existence. Instead, the focus of the field has shifted to the development of more generalizable epitope-agnostic models which are designed to predict the binding of any epitope and TCR by learning the epitope-TCR interaction patterns from a limited set of epitopes. The latest IMMREP benchmark (IMMREP23) demonstrated that epitope-agnostic models still behave like random predictors for unseen epitopes, i.e., epitopes that are not present in the training data (9). A possible explanation is the presence of insufficient high-quality training data (10). However, to date, it is unknown how much training data is required to achieve acceptable performance.

Although, initially, most focus was given to the development of new methods achieving better predictions, the latest research emphasizes the importance of data quality and the influence of biases in training data on model performance. Non-biological biases can have a high impact on the prediction model resulting in unrealistic performances. For example, Luu et al. highlighted the importance of consistent data processing, i.e. all training and test data sets must undergo the same processing steps (10). Currently, different databases are storing epitope-specific TCR data, each having their own criteria of data parsing and quality control (10). Slight variations in the notation of the TCR or epitope sequences in the training data could be seen as epitope-specific by the model and thus affect the predictions resulting in an inaccurate model performance (10). Another important aspect when considering training epitope-TCR recognition models, is the generation of the negative training data set (11,12). In theory, the negative training data must consist of TCRs not binding the epitope of interest. However, no experimental procedures exist to determine representative non-binding TCRs accurately in a high-throughput manner (11). Hence, alternative options were explored to construct negative training datasets. These include the random selection of TCRs from non-stimulated TCR repertoire data and shuffling epitope-TCR pairs within the positive training data (10,11,13). For both strategies, the absence of epitope-specific TCRs cannot be guaranteed due to the presence of epitope-specific TCRs in naïve repertoires (14,15) and the phenomenon of cross-reactivity resulting in TCRs binding more than one epitope. However, in the case of epitope-agnostic models, the shuffling method has been found to be the lesser of evils as this avoids the presence of distribution biases in the TCR or epitope sequences (11).

To fully understand the success and limitations of current epitope-TCR prediction models, it is essential to investigate the sequence distribution of the training data and its connection with the model performance. In this paper, we define sequence distribution as the mapping of the TCR β amino acid sequences into a latent space, enabling the study of the underlying overlap in TCR sequences recognizing the same epitope. We refer to the positive training data as epitope-specific TCR repertoires. In short, an epitope-specific TCR repertoire is a collection of TCRs all targeting the same epitope. In contrast to the typical TCR repertoires, these epitope-specific repertoires contain TCRs from different individuals collected over different studies. It is well established that a single epitope presented by a specific MHC molecule can be recognized by highly diverse TCR sequences (16). In general, epitope-specific TCR repertoires contain multiple TCR groups with different amino acid patterns or motifs (10). Previous research has already demonstrated a link between the epitope-specific TCR diversity and model performance, where a more diverse epitope-specific TCR space, and therefore training data set, often results in less performant prediction models (4,5,17,18). In this paper, we sought to understand the general sequence composition within epitope-specific TCR β repertoires by performing an in-depth study on a common training data set. To this end, we used a cluster-based approach to enumerate the sequence diversity of unique TCR repertoires and simulated epitope-specific TCR repertoires to study the origin of this diversity. The found associations between TCR data composition and model performance provide new insights into the correct interpretation of performance curves when validating models and help our understanding of the optimal datasets required to train well-performing epitope prediction models.

## Materials and methods

### Data collection

To understand the general sequence distribution of epitope-specific TCR β repertoires, a large collection of epitope-specific TCR β data was needed. Since the current study focuses on the relationship between different metrics derived from epitope-specific TCR repertoires and the performances of their prediction models, the TCRex model training data was used (7). This includes a combined data set from several public databases, including VDJdb (19), McPAS-TCR (20) and the ImmuneCODE™ database (21). Every TCR β chain (i.e., unique combination of V/J gene and CDR3 β sequence) is included only once per epitope to reduce overrepresentation of TCR signatures by duplicates. Both positive and negative TCRex training data was used in this study. All data was parsed according to TCRex version v0.3.3 standards. In total, TCR β chain data was collected for 6 cancer and 120 viral epitopes.

### Prediction model

The TCRex prediction model was used as the performance baseline for this study as it has several properties that can be easily leveraged toward a broader understanding (7). Firstly, it is highly performant as per the IMMREP22 benchmark (22) since it outperforms more complex methods at single-epitope classification. Secondly, it is built on random forest classifiers, which allows some level of model explainability. Thirdly, it is fast in both training and application, allowing for maximum tractability on larger and variable data sets. Training data was included if at least 30 different TCRs per epitope were available. The TCRex tool automatically makes a distinction between acceptable and not acceptable models. The latter includes models with the area under the receiver operating characteristic (ROC) below 0.7 and/or an average precision below 0.35 and thus not accessible at the online TCRex platform. In addition to positive data, the typical TCRex workflow randomly collects negative data from a large negative data set containing TCR sequences derived from different healthy individuals (23). This is common practice for single-epitope models as they are intended for use in screening full repertoires for epitope-specific TCRs. In general, 10 times more negative TCRs are used than epitope-specific TCRs when training prediction models. However, if negative TCRs share the same CDR3 β sequence and V/J genes as an epitope-specific TCR, they are excluded from the training data set.

### Clustering the entire collection of epitope-specific TCR sequences

Previous studies have already illustrated the presence of different clusters of TCRs within epitope-specific repertoires (24,25). However, it was not yet clear how TCR clusters differed between epitopes. To assess whether TCR clusters of the same epitope were more similar in their CDR3 β sequence than clusters of different epitopes, all collected epitope-specific TCRs were clustered using a Hamming distance of one on the amino acid level with clusTCR (26). Given the limited size of the dataset (< 50 000 TCRs), the single-step Markov Cluster (MCL) approach was used. Clustering was done on the CDR3 β level only as we wanted to collect similar CDR3 sequences with different V genes in the same cluster. The results were visualized as graphs with NetworkX (27).

### Evaluating epitope-specific TCR motifs across epitopes

In order to study the possible overlap in TCR β sequences recognizing different epitopes, epitope-specific motifs were generated from the positive training data sets, i.e. epitope-specific TCR repertoires, and subsequently clustered by clusTCR. Motif generation itself also included a clustering step: for every epitope, all collected specific TCRs were clustered based on their CDR3 β sequencing and a hamming distance of one using the MCL setting in clusTCR. This resulted in a list of clusters containing two or more CDR3 β sequences. ClusTCR automatically generates a motif for each cluster summarizing its CDR3 β content. These motifs are consensus sequences showing one or more amino acids per position depending on their frequencies at the respective position. If one amino acid is dominating a position, i.e., being present in at least 70% of the sequences in the cluster, it will be written in upper case in the final motif. If two amino acids are dominating a position almost equally, both are written in upper cases between square brackets. If they differ largely in frequency, only the most dominant amino acid is selected, but it is written in lowercase. In case of complete promiscuity at a position, a dot is written (26). After clustering every epitope-specific repertoire separately, a list of sequence motifs was collected per epitope. To evaluate if motifs associated with the same epitopes are more similar that motifs derived from other epitope-specific TCR repertoires all motifs were clustered again using the same approach. However, since clusTCR can only cluster sequences containing A-Z characters (e.g. amino acids), all non-amino acid characters and lowercase characters in the motifs were first replaced by character X, indicating variability at that position.

### Visual representation of the TCR space with UMAP

To visualize the similarity in TCR sequences across different groups (e.g. epitopes), RapTCR was used (28). In brief, this tool creates a two-dimensional representation of the TCR similarity space, in two steps. First, each CDR3 sequence is embedded as a vector of fixed length, where the Euclidean distance between vectors approximates their similarity as defined by TCRdist (5). Next, the dimensionality of this vector set is reduced by applying Uniform Manifold Approximation and Projection (UMAP) (29), enabling two-dimensional visualisation. All UMAPs were created with the number of neighbors set to 15, and the minimal distance to 0.8.

### Simulation of epitope-specific TCR repertoires

Synthetic TCR repertoires were generated *in silico* using the rejection sampling strategy within the LIgO AIRR simulation software suite (30). Briefly, the rejection sampling method generates a large number of TCRs and then retains those that match a user-defined motif. The motifs were defined by a seed, i.e. a short amino acid sequence, and a range of allowed hamming distances. To create realistic motifs, the seeds were derived manually from epitope-specific TCRs stored in the VDJdb, by removing the first and last three amino acids from the CDR3 β sequences (Supplemental Material S1). Due to the selection of longer seeds, a hamming distance of up to three was allowed. 300 TCRs were generated per motif. In total eight synthetic TCR repertoires were generated, each starting from three LIgO seeds. The YAML files with all parameters and instructions defining LIgO simulations can be found in the GitHub repository. The final simulated repertoires were clustered using clusTCR.

### Generating new TCRs to fill the gaps in between single TCRs and clusters

Clustering epitope-specific TCR data most often results in clusters of different sizes and unclustered TCRs. We refer to the latter as singlets. To understand the origin of these clusters and the absence of one large cluster, we introduced new TCR sequences that fit in between these clusters and linked them together. This was done in two separate studies using a similar strategy: (1) clustering clusters together and (2) clustering singlets with another singlet or cluster (Figure 1 B). For simplicity, we explain the method that was used to link a singlet with another TCR or cluster. To avoid clustering based on TCR length, the hamming distance of one was replaced with a Levensthein distance of one. For every singlet, we generated a list of new TCRs, so that we can link it with at least one other singlet or cluster. To this end, we calculated the Levensthein distance between the singlet and any other TCR within the data set and selected those TCRs with the shortest Levensthein distance. (In the case of generating TCRs in between clusters, the same strategy was used for every TCR in a cluster to find all TCRs with the shortest Levensthein distance that are not present within the same cluster.) For every of these selected TCRs, we selected the best global alignment with the singlet using the biopython (31) pairwise2.align.globalms function with a match score of 2, a mismatch score of -1, a gap opening penalty of 1.1 and a gap extension penalty of 0.6. New TCRs were generated iteratively, by first replacing the gaps in one of the sequences with the matching amino acid present in the other. After every sequence-update, the two sequences were realigned and the process was repeated. When all gaps were resolved, the same process was repeated to fix the mismatches. One mismatch was allowed in the final alignment as this corresponds with a Levensthein distance of one. The final list of TCRs that link the singlet with another TCR in the data set is given by all the TCRs updated during the alignment iterations. To study the generation probability of the clustered versus the linking TCRs, a new set of 8 TCR datasets were simulated. These were derived by generating 3000 TCRs for the first seed of every simulated repertoire. Thus, 8 new TCR datasets were simulated, each containing 3000 TCRs derived from one seed. The same analysis was also performed on the 8 epitope-specific TCR repertoires in our TCRex database containing the most TCR data. Before generating the new TCRs for these simulated and experimental repertoires, all TCRs with a generation probability of zero were removed.

**Figure 1:**
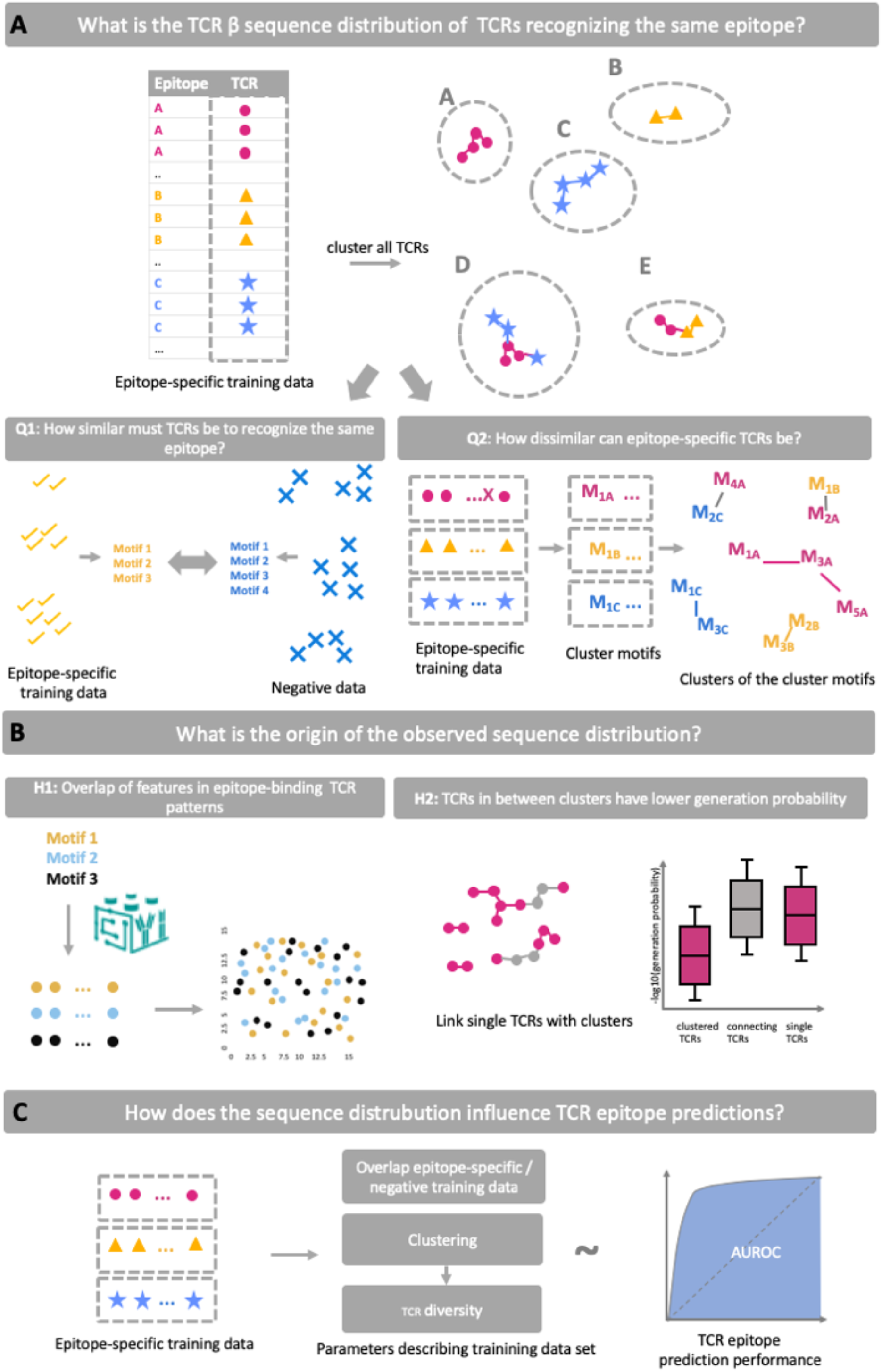
Graphical summary of the main research questions and methods. **(A)** This study starts with an exploration of the TCR β sequence distribution in epitope-specific TCR repertoires by clustering all collected TCR sequences for all epitopes at once. The resulting clusters are either pure, i.e., all TCRs are specific for one epitope (e.g. clusters A, B and C), or impure when TCRs against multiple epitopes are grouped together (e.g. clusters D and E). Consequently, TCRs recognizing the same epitope can share similar sequences and belong to the same cluster or share dissimilar sequences and belong to different clusters. In the latter case, TCRs sequences recognizing different epitopes might even share more similarities than TCRs recognizing the same epitope. These initial results introduce two different questions: (1) “How similar must TCR sequences be to recognize the same epitope” and (2) “How dissimilar can epitope-specific TCR sequences be?” To answer the first question, the epitope-specific TCRs motifs were compared with motifs hidden in the negative training data. The second question was tackled by clustering every epitope-specific TCR repertoire separately to generate epitope-specific motifs, which were clustered in a second step. **(B)** After an initial investigation of the TCR β sequence distribution, we aim to understand the origin of this distribution. First, we hypothesized that TCRs recognizing different epitopes can have overlapping epitope-specific patterns. To test this hypothesis, multiple TCR repertoires were simulated with LIgO using specified motifs and the sequence distribution was evaluated across these motifs. Second, instead of one big cluster per epitope, various clusters per epitope-TCR specific repertoire can be found. We hypothesize that all these TCRs belong to a single cluster, but due to the lack of observation of TCRs with lower generation probability in the data, this cluster has been separated into multiple smaller clusters. To investigate this hypothesis, we compared the generation probabilities between (un) clustered TCRs and TCRs linking the observed clusters or single TCRs together. For simplicity only the connection of single TCRs to other clusters is shown in the figure. **(C)** Finally, the effect of the sequence distribution on TCR-epitope prediction was assessed. Three different metrics were calculated per epitope-specific TCR repertoire: the overlap in CDR3 β sequence with the negative training data, the percentage of clustered TCRs and the TCR diversity. Each of these metrics were compared with the area under the receiver operating characteristic curve (AUROC).

### Cluster-based diversity metrics to enumerate unique TCRs

Different metrics exist to summarize the diversity of TCR data sets (32).However, these are based on the presence of TCR clones having different clone counts. In our epitope-specific data sets however, every TCR (V/J gene + CDR3 β) is included only once. Hence, a cluster-based diversity metric is calculated based on the number and size of the clusters in the data set. In short, every epitope-specific TCR data set is clustered separately using clusTCR. Hereafter, the size of the clusters is calculated and used as a proxy for clone counts, while the clusters are seen as TCR clones. Singlets, i.e. TCRs not belonging to a cluster of at least 2 TCRs, are considered a clone with only one count. Thus, traditional diversity metrics can be used on these clustered data sets. Here, the Shannon diversity index, Gini-Simpson index, Pielou’s index and Gini-coefficient were calculated for each epitope-specific TCR data set using scikit-bio (33). In addition, the Diversity Evenness 50 (DE50) value was calculated as demonstrated in (32).

## Results

### Overview of the collected data

Epitope-specific TCR β chain data was collected for 6 cancer and 120 viral epitopes. Due to several large initiatives during the COVID-19 pandemic, the majority of the collected TCR data covers SARS-CoV-2 epitopes. To evaluate the predictive capacity of each epitope towards unseen TCR sequences, we trained TCRex prediction models in a typical cross-validation setup. For 98 epitopes, the model demonstrated a good performance on held-out TCR sequences (Supplemental Material S2). The remaining epitope models, 28 in total, failed to achieve an acceptable performance as per the defined criteria, i.e. AUROC below 0.7 and average precision below 0.35. A more thorough overview of all training data and the performances of the resulting models can be found in Supplemental Material S2.

### Epitope-specific TCR repertoires are an ensemble of different TCR clusters

To enumerate the potential diversity of TCR β sequence motifs, groups of similar TCRs were quantified in the collected data set of 42 050 unique epitope-specific CDR3 β sequences. 34% of the TCRs, i.e. 14215 TCRs) could be clustered using a Hamming distance of one, resulting in 2 121 clusters. Clusters varied in size from 2 TCRs up to 319 TCRs (Supplemental Material S3). Two types of clusters can be observed: pure clusters (i.e., containing CDR3 β sequences associated with only one epitope) and non-pure clusters of CDR3 sequences associated with multiple different epitopes. Of the 2 121 clusters, 796 are pure. These clusters are small ranging from 2 up to 36 sequences (Supplemental Material S3).

### Epitope-specific TCR clusters are spread out over TCR sequence space

Since 62% of the clusters were not pure, we hypothesized that CDR3 β sequences share motifs across epitopes. To investigate cross-epitope motif sharing, every epitope-specific TCR repertoire was summarized as a list of CDR3 β motifs, which represent the general TCR signatures of every epitope-specific cluster in the repertoire. In contrast to the previous clustering strategy, every epitope-specific repertoire was thus clustered separately to obtain epitope-specific motifs, resulting in a total of 1921 unique motifs. Hereafter, all generated motifs were clustered together to evaluate whether TCR clusters sharing the same epitope specificity were more similar than clusters with other specificities. ClusTCR clustered 464 of 1921 motifs in 136 clusters: 29 pure and 107 impure clusters (Supplemental materials S4). In total, 56 motifs were found to be shared between different epitopes. These epitopes can have both a very similar and dissimilar protein sequence (Supplemental materials S4). These clustering results support the idea that epitope-specific TCRs are spread out over TCR sequence space, i.e. TCRs with entirely different epitope specificity might sometimes be more similar in their amino acid sequence than TCRs recognizing the same epitope. The same conclusion can be visually supported by a UMAP representation of the TCR similarity space (figure 2). Here, every point represents a TCR sequence. For visualization purposes, only three epitopes were selected to represent its epitope-specific TCRs. We observed that every color is spread across the entire UMAP space, signifying that TCRs with identical epitope-specificity can differ in terms of amino acid composition.

**Figure 2:**
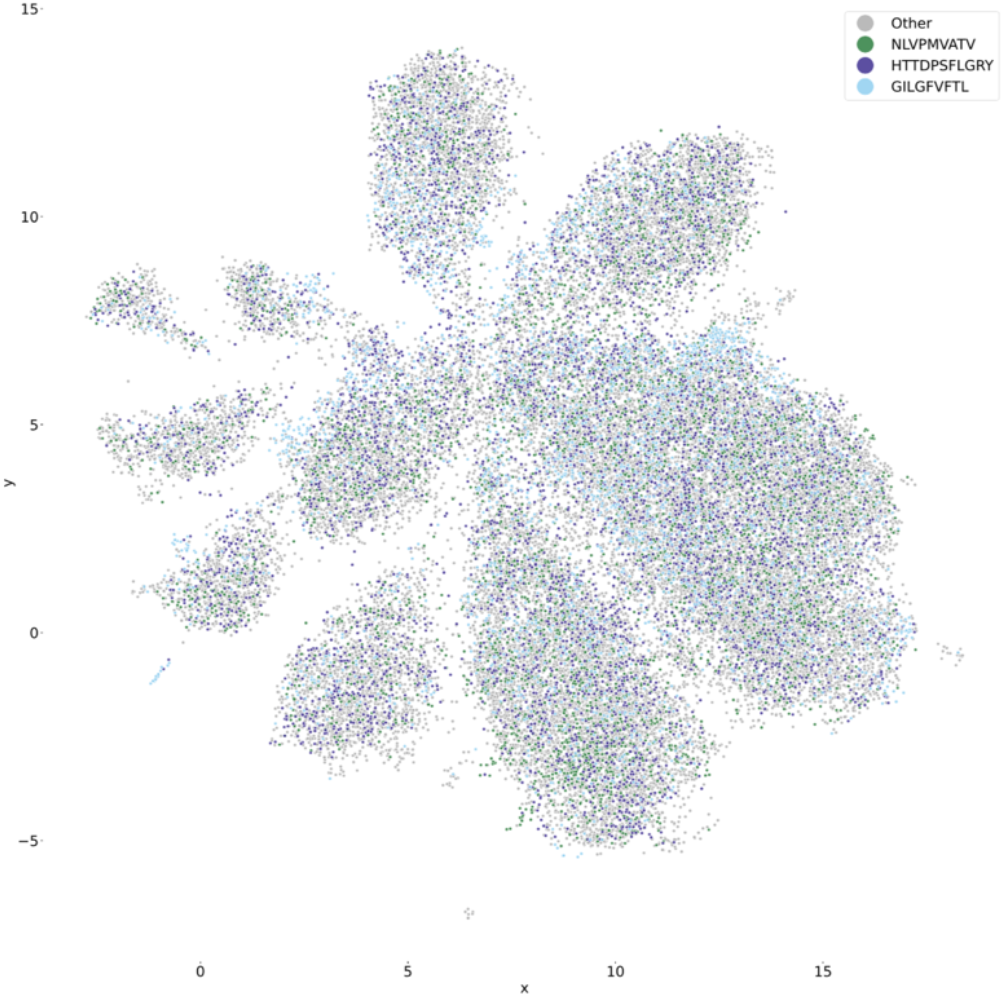
Epitope-specific TCRs are spread out over TCR sequence space. UMAP of epitope-specific TCRs colored by epitope-specificity. The gray dots represent TCRs that are not linked solely with one of the three selected epitopes, i.e. they can bind other epitopes or one or more of the selected epitopes and other epitopes.

### Negative TCRs can differ in only one amino acid with epitope-specific TCRs

Training an epitope-specific TCR prediction model requires both TCRs known to bind the specific epitope, hereinafter referred to as the epitope-specific TCR data, and TCRs that are not expected to bind the epitope, i.e. the negative training data set. Since one amino acid difference might affect the affinity of a TCR for an MHC-presented epitope, some non-epitope-specific TCRs might look very similar to epitope-specific TCRs. To investigate the overlap between epitope-specific and negative TCRs, non-shared epitope-specific and negative training TCRs were clustered for each epitope separately. As shown in Supplemental materials S5, 19% of all clusters contain a mix of both epitope-specific and negative data. Hence, some CDR3 β sequences in the negative data set differ with only one amino acid with the epitope-specific TCRs. Indeed, when studying the minimal distance between every TCR in the epitope-specific TCR dataset and its closest negative TCR (Supplemental Material S5), minimal Levenshtein distances ranged from 0 until 3. TCRs with a minimal Levenshtein distance of 0 contain identical CDR3 β sequences in the epitope-specific and negative datasets. They do however differ in their V/J genes as exact overlap between epitope-specific and negative training data was avoided during creation of the training data. These results raise the question of whether clusters with similar motifs are shared between the epitope-specific data and negative data. To investigate this, the epitope-specific and negative data were clustered separately using the same clustering method as before. Motifs were compared between epitope-specific and negative clusters. To avoid the generation of clusters due to shared CDR3 β sequences between the two data sets, these sequences were removed entirely before clustering. For seven epitopes, identical motifs were found between epitope-specific and negative clusters (Supplemental material S5). An example is shown for the training data of epitope HTTDPSFLGRY in figure 3. All negative data for this SARS-CoV-2 epitope was collected before the COVID-19 pandemic. The results show that the negative TCRs are scattered throughout the epitope-specific TCR space, i.e. epitope-specific TCRs can be more similar to negative TCRs than other epitope-specific TCRs.

**Figure 3:**
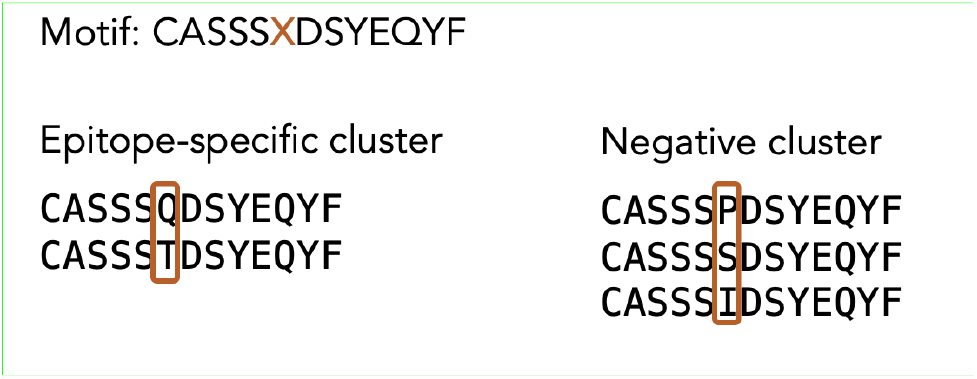
Example of clusters in the epitope-specific and negative training data sharing a sequence motif. Both clusters were derived from the training data of epitope HTTDPSFLGRY by clustering the epitope-specific TCRs and negative TCRs separately. The five TCR sequences differ in only one position. As a result, the two clusters share the same consensus motif.

### TCRs originating from different LIgO seeds can have similar sequences

Recently, LIgO was released to enable the simulation of TCR or BCR receptor and repertoire data (30). The simulation of TCR sequences from defined seeds facilitates the generation of large TCR data sets with preset TCR patterns, overcoming current limitations in the public epitope-specific TCR data space, such as the limited number of TCRs sharing a motif (34). To study the potential origin of the observed sequence distribution of epitope-specific TCR repertoires, eight repertoires were simulated with LIgO (30) based on previously identified epitope-specific TCR sequences. Each repertoire was generated using three LIgO seeds derived from experimental TCRs recognizing the same epitope, i.e. FLKEKGGL, GTSGSPIVNR, YVLDHLIVV, LLWNGPMAV, KTFPPTEPK or YSEHPTFTSQY (Supplemental materials S1).

Although only three LIgO seeds were used to simulate every epitope-specific TCR repertoire, clustering the resulting synthetic repertoires results in a multitude of clusters each with their own specific consensus motif (Supplemental materials S6). Most of these clusters group TCRs originating from the same LIgO seed. However, some clusters contain TCRs with a different LIgO seed origin within one simulation, showcasing that TCRs containing different binding motifs can still be very similar to each other (Supplemental materials S6). Indeed, in the UMAP visualizations (figure 4, Supplemental materials S7) we can see a natural grouping between TCRs that originated from very different LIgO seeds. Finally, clustering the consensus motifs from the different simulated repertoires, each having a different seed origin, shows that some cluster motifs are more similar to cluster motifs derived from other LIgO seeds than from the same LIgO seeds (Table 1). Two motifs are even shared by two clusters from different simulations due to the presence of identical CDR3 β sequences in both clusters (Supplemental materials S6). This is likely a result of similarities within their seeds of origin, i.e. SHSLAGGSNE (simulation 6) and SPLLAGGPYE (simulation 8). Thus, the clustered diversity of our epitope-specific TCR repertoire might be explained by the overlap of features in epitope-binding motifs.

**Table 1:**
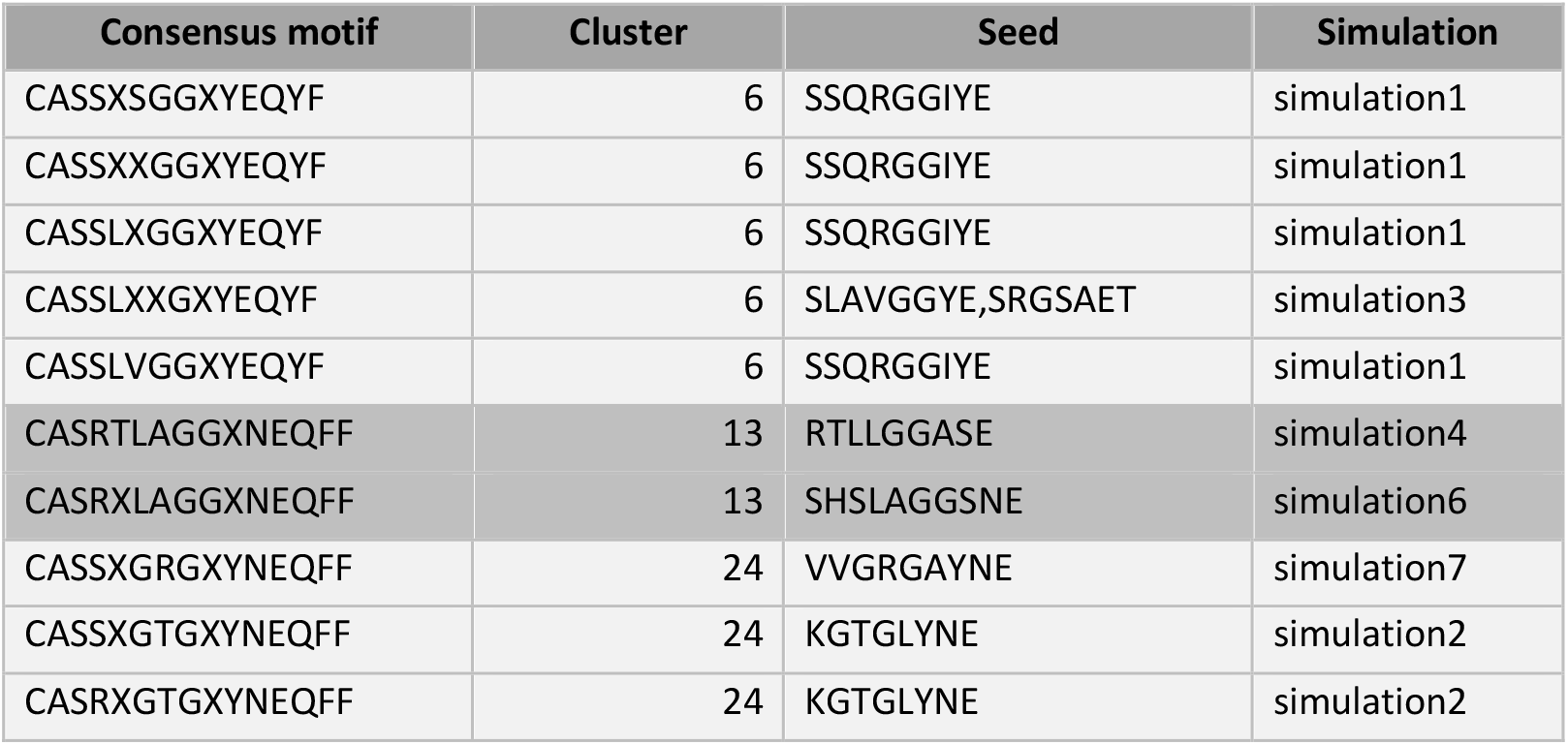
Overview of the clusters grouping consensus motifs derived from different simulations. Every cluster groups two or more consensus motifs. For every of these motifs, the LIgO seed is shown that gave rise to the TCR sequences within the original clusters making up these consensus motifs.

| Consensus motif | Cluster | Seed | Simulation |
| --- | --- | --- | --- |
| CASSXSGGXIEQYF | 6 | SSQRGGIYE | simulation1 |
| CASSXXGGXIEQYF | 6 | SSQRGGIYE | simulation1 |
| CASSLXGGXIEQYF | 6 | SSQRGGIYE | simulation1 |
| CASSLXXGXIEQYF | 6 | SLAVGGYE,SRGSAET | simulation3 |
| CASSLVGGXIEQYF | 6 | SSQRGGIYE | simulation1 |
| CASRTLGGXNEQFF | 13 | RTLLGGASE | simulation4 |
| CASRXLAGGXNEQFF | 13 | SHSLAGGSNE | simulation6 |
| CASSXGRGXNEQFF | 24 | VVGRGAYNE | simulation7 |
| CASSXGTGXNEQFF | 24 | KGTGLYNE | simulation2 |
| CASRXGTGXNEQFF | 24 | KGTGLYNE | simulation2 |

**Figure 4:**
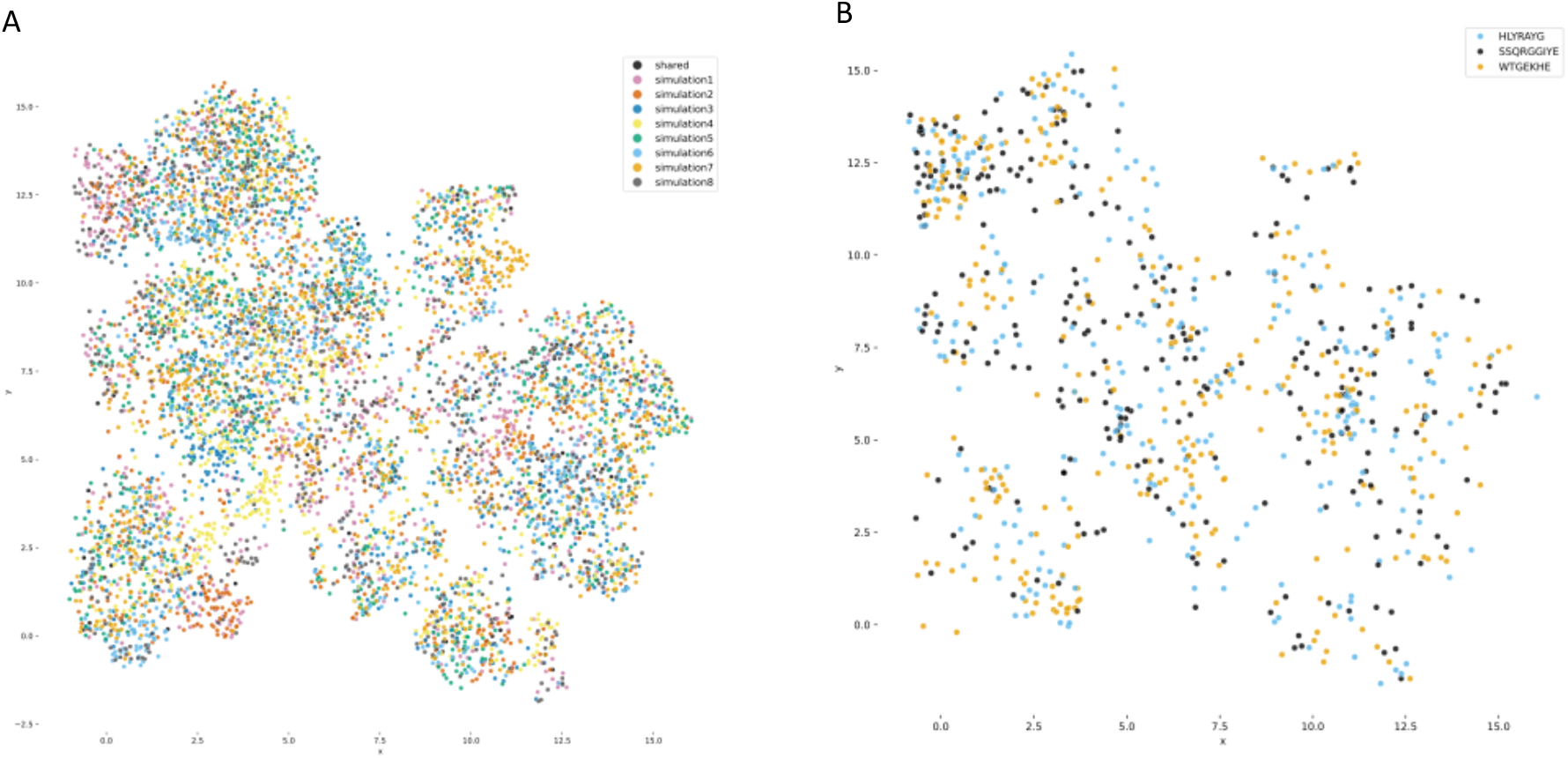
Spread of simulated TCR repertoires in space, where each simulation is a proxy for a single epitope with three ground-truth TCR motifs. **(A)** UMAP plot of all LIgO-simulated repertoires colored by simulation, i.e. simulation 1 up to simulation 8. The 96 TCRs that were simulated by more than one simulation are colored black. **(B)** UMAP plot of simulation 1, colored by seed. The shared TCR sequences were not shown.

### TCRs in between clusters have low generation probability

In the previous section, we focused on the presence of dissimilar TCRs within epitope-specific TCRs repertoires sharing different motifs, thus resulting in different clusters. However, not all epitope-specific clusters carry distinct motifs. Often, multiple clusters with highly similar motifs can be observed, i.e., TCRs sharing the same sequence motif can still be grouped in separate clusters. We hypothesize that this is the result of a difference in generation probability where TCRs linking clusters together have a lower generation probability and are thus less likely to be present in any repertoire. Therefore, TCRs with the same sequence motif would be in the same cluster if it were not for gaps in the TCR space due to these being more difficult to generate. To study this hypothesis, we generated TCRs that link two clusters or a single TCR with a cluster for the eight TCRex-repertoires containing the most TCR data. Figure 5 demonstrates that these linking TCRs have a lower generation probability compared to clustered TCRs, which is the reason that we are unlikely to observe them resulting in multiple smaller clusters instead of one big cluster per motif.

**Figure 5:**
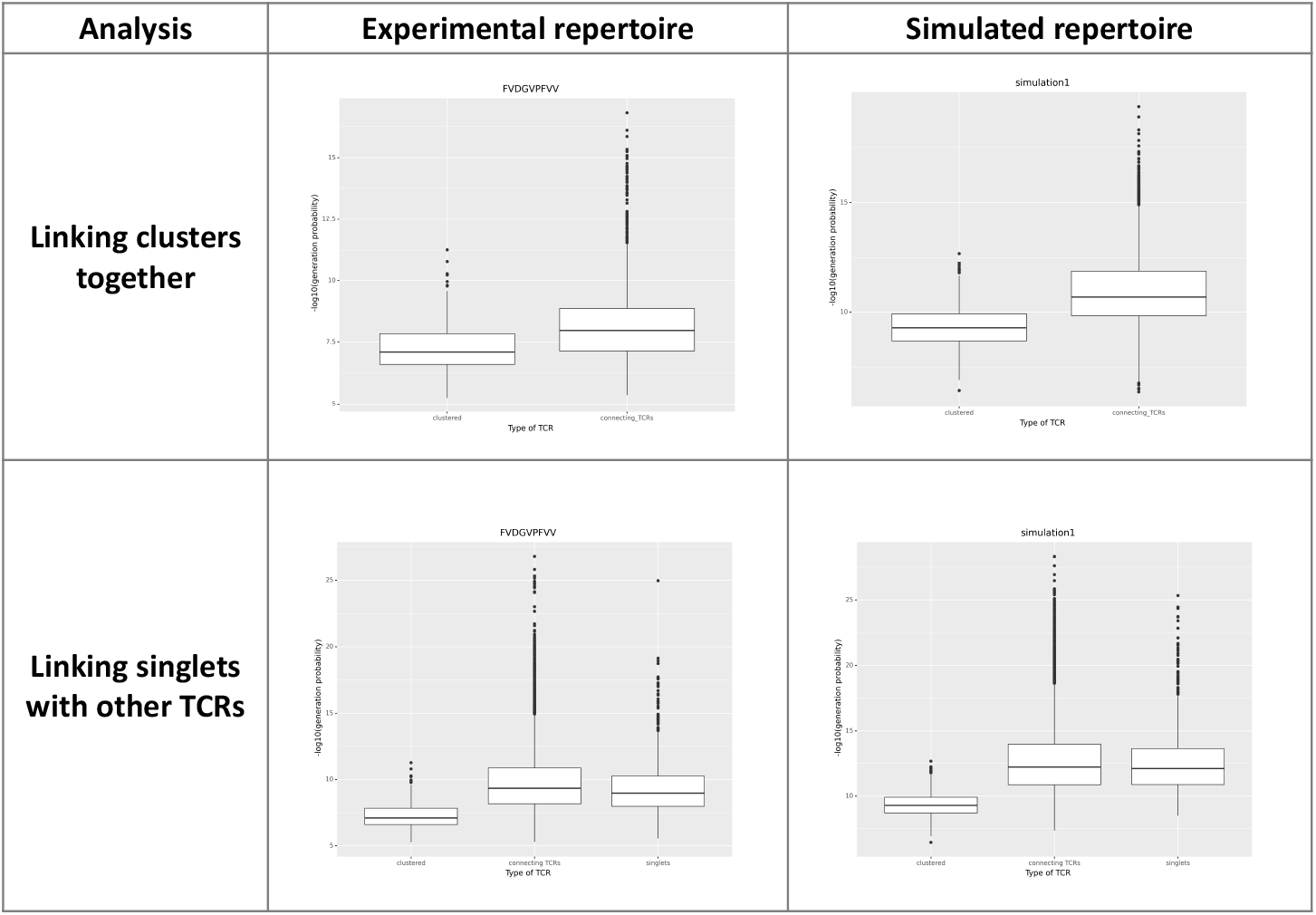
Generation probability distribution for clustered TCRs, singlets and new linking TCRs. For each of the 8 selected experimental repertoires and 8 simulated repertoires, the generation probability was calculated for all TCRs within the repertoires and those that were newly generated to link clusters or singlets to other TCRs in the data set. Here, one experimental and one simulated repertoire is selected to demonstrate the differences in generation probability. The negative of the common logarithm of the generation probabilities are plotted on the y-axis. Hence, the lower the generation probability, the higher the value on the y-axis. The plots for the remaining epitopes and simulation show similar results and are reported in Supplemental material S8.

The study of the generation probability distribution on experimental (i.e., present in the TCRex database) epitope-specific TCRs holds a few limitations. First of all, the number of TCRs per sequence motif is limited to those that have been previously identified. Thus, the absence of TCRs connecting the different clusters might be a result of limited sampling instead of a very low generation probability. Secondly, the exact TCR generation model is unknown for the sampled individuals, introducing possible small errors when calculating generation probabilities according to one “average” IGoR model. Both limitations can be circumvented by generating repertoires *in silico*: the number of TCRs simulated per motif can be increased and it is guaranteed that you simulate and evaluate each sequence with exactly the same model. Hence, we investigated the distribution of the generation probability on LIgO-simulated TCR data sets. Every data set contains approximately 3000 TCR β chains generated from only one seed, i.e., the first seed that was used to generate the original simulated data set with three seeds. These one-seed based simulated TCR repertoires were then clustered separately using a Levenshtein distance of one. Figure 5 and Supplemental material S8 show a similar relationship between the generation probabilities of TCRs for simulated repertoires and experimental repertoires, thus supporting our hypothesis that ‘gaps’ observed in between epitope-specific TCR repertoires are a result of TCR sequences with a lower generation probability.

### Large overlap between epitope-specific and negative TCR sequences influences model performance

Previous analyses showed that there can be a substantial overlap between the epitope-specific and negative data set of an epitope-specific model, i.e., similar sequences can be present in both. This might have an influence on the final performance of the trained model, as a larger overlap reduces the separation of epitope-specific and negative TCRs in space. To assess the possible relation between training data overlap and model performance, the minimal Levenshtein distance was calculated for each CDR3 β sequence between its closest epitope-specific and negative TCR partner. Hereafter, the minimal positive distance was divided by the minimal negative distance for every sequence. For each model, the median of these fractions was calculated resulting in one number representing the minimal distances in the training data set. As shown in figure 6, models trained on data with high overlap generally show a worse (AUROC) performance.

**Figure 6:**
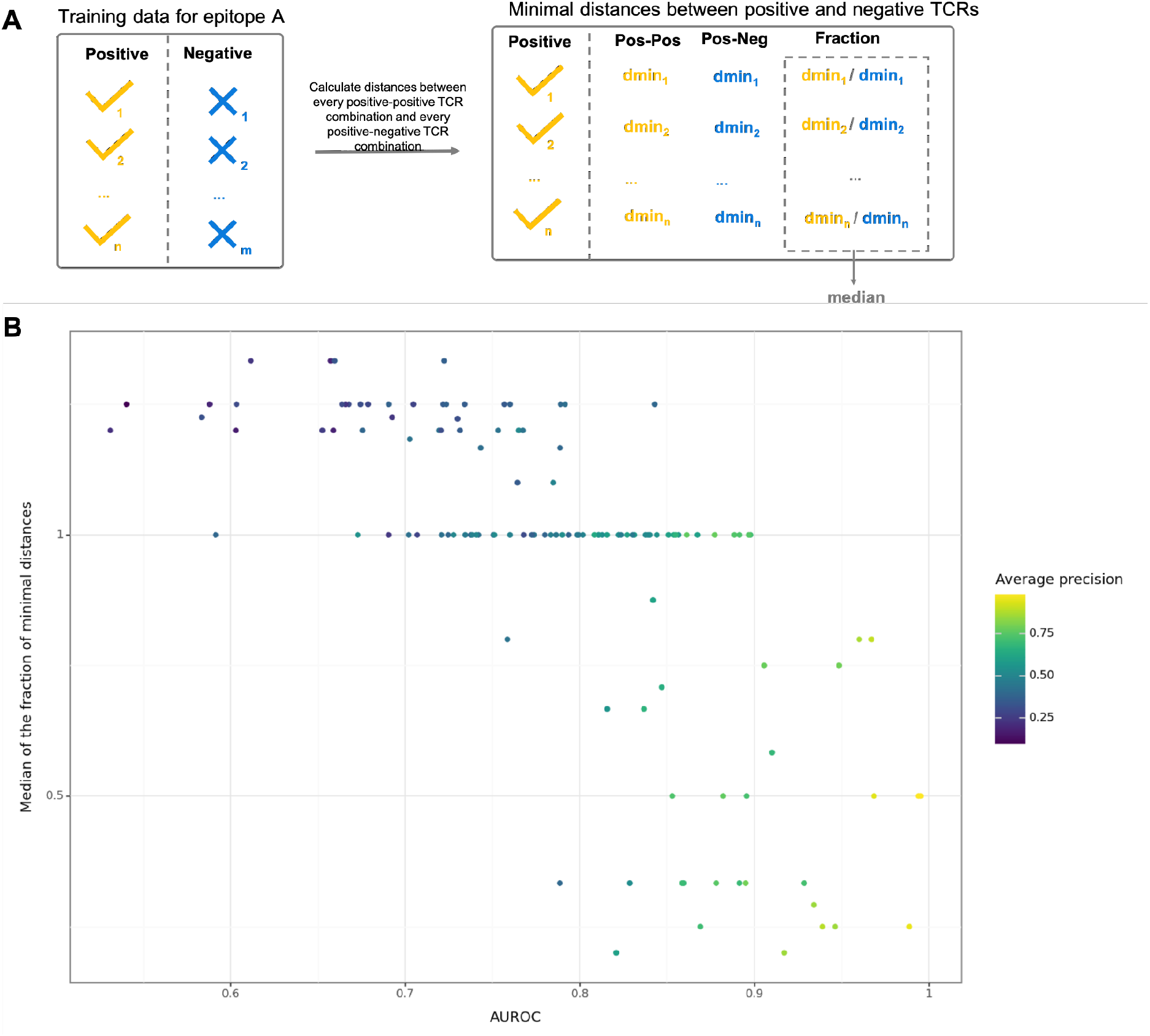
Models trained on data with high similarity between the epitope-specific and negative TCR sequences show poorer performance. **(A)** Graphical overview of the method to calculate the median of the fraction of minimal distances. For every TCR in the epitope-specific training data set for epitope A: (1) the levenshtein distance is calculated with all other epitope-specific TCRs and all negative TCRs, (2) the minimal distance is selected for both groups (3) the minimal distance with an epitope-specific TCR is divided by the minimal distance with a negative TCR. This results in a list of fractions per epitope. **(B)** The median of the fractions of minimal distances between epitope-specific and negative training data versus AUROC is given. A number higher than 1 represents a model where the median epitope-specific TCR is more similar in its CDR3 β sequence with negative sequences than other epitope-specific TCRs. In general, higher AUROC values are seen when the minimal distances are smaller within epitope-specific TCR repertoires than in between these repertoires and the negative training data.

### Model performance is linked with the number of TCRs that can be clustered within the epitope-specific training data

A high overlap in TCR signatures within the epitope-specific training data is expected to make the identification of epitope-specific signatures easier and thus result in a good model performance. Hence, we expected a clear relationship between the model performance and CDR3 β sequence overlap in the epitope-specific training data. To test this hypothesis, each epitope-specific TCR data set was clustered and the number of clustered TCRs was plotted together with the AUROC of its epitope-specific model. A clear trend is seen between the percentage of clustered TCRs within the epitope-specific data and the model performance (figure 7A).

**Figure 7:**
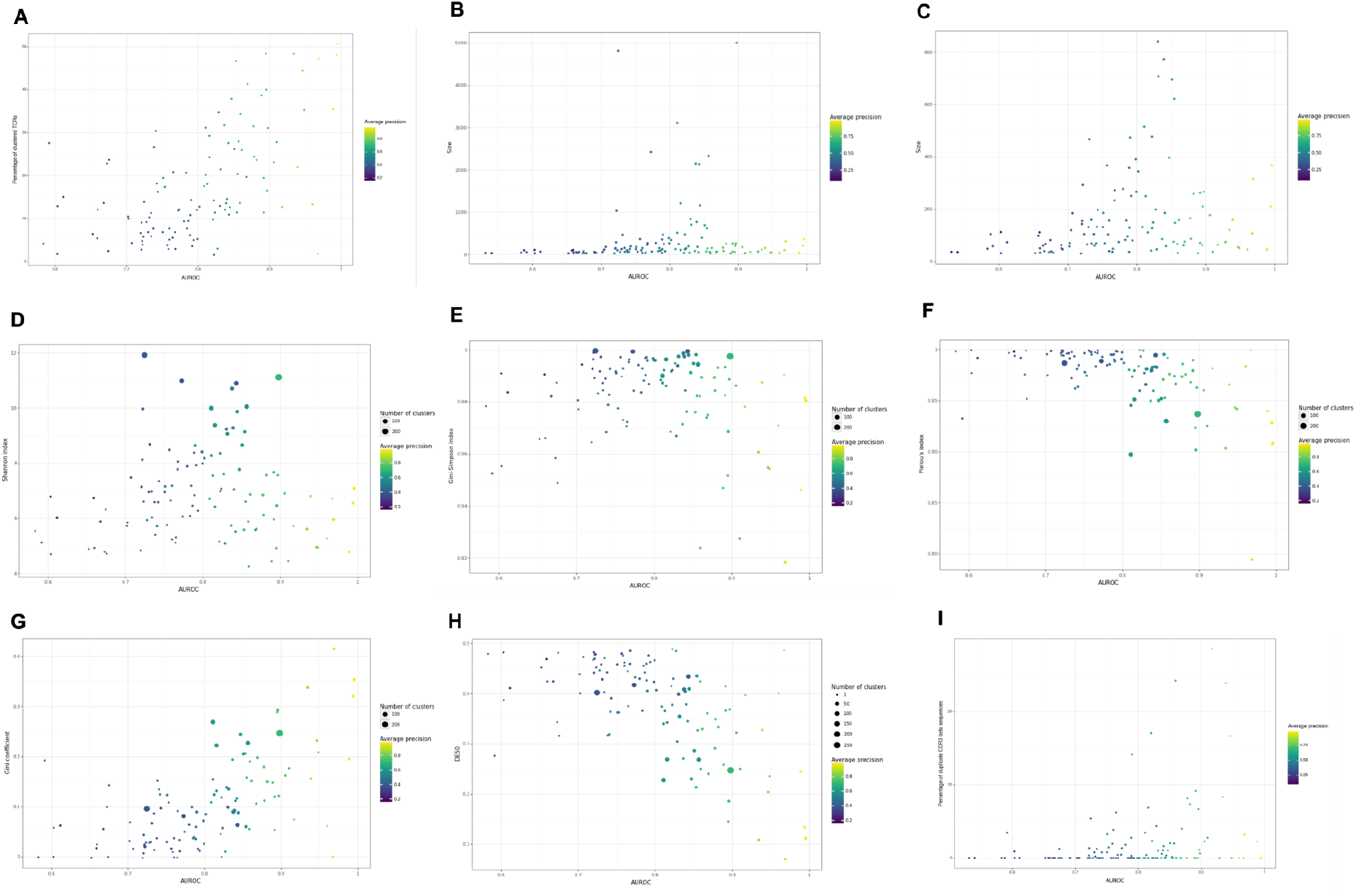
Relation between the cluster capacity, size and diversity of the epitope-specific TCR training data set and model performance. **(A)** Percentage of clustered epitope-specific TCRs versus AUROC, colored by average precision **(B**,**C)** Relation between the size of the epitope-specific TCR training data set and model performance for all epitopes **(B)** and all epitopes having less than 1000 epitope-specific TCRs **(C)**. Relation between diversity and model performance as expressed by the **(D)** Shannon index, **(E)** Gini-Simpson index, **(F)** Pielou’s index, **(G)** Gini coefficient and **(H)** DE50. The size of the dots refers to the number of clusters in each clustered epitope-specific TCR repertoire. Figure **(I)** represents the percentage of duplicate CDR3 β sequences in the training data versus the AUROC.

### Clonal epitope-specific training data sets result in more performant models

In general, a larger training data set is expected to result in a better performing model. However, figures 7B and 7C demonstrate the absence of a clear relationship between the size of the training data and the model performance. We hypothesized that the TCR diversity of the training data has a large influence on the model performance and thus large data sets with large TCR diversity might result in lower model performance in comparison with models trained on smaller training datasets having less diversity within their TCR sequences. To investigate the relation between TCR diversity and model performance each epitope-specific TCR data set was clustered separately using clusTCR and the diversity of the clustered data sets were described using five different metrics. The Shannon diversity index and Gini-Simpson index are often used for the analysis of full TCR repertoires. In figure 7D and 7E no clear relationship can be found between these diversity metrics and the AUROC. When comparing figure 7D with figure 7C, the Shannon diversity seems to be largely affected by the size of the epitope-specific TCR repertoires, which is a common feature of this diversity metric. The Pielou’s index is a normalized Shannon index and thus gives a better representation of the diversity over different repertoire sizes. Figure 7F shows a drop in the Pielou’s index with increasing AUROC values indicating a relationship between the evenness in cluster sizes and model performance. The same conclusion can be drawn from Figure 7G where the higher values for the Gini coefficient are associated with higher AUROC values and Figure 7H where lower DE50 values are associated with higher AUROC values. Thus, training data sets containing more clonal epitope-specific TCR repertoires result in better performing models, suggesting that the presence of a few large clusters drive the model performance. Although clustering was done on a list of unique CDR3 β sequences, some training data sets do contain multiple instances of identical CDR3 β sequences with different V/J genes. The presence of these replicates can also influence the model performance as these replicates could be present in both training and testing data sets during cross validation (figure 7I).

## Discussion

Training a decent prediction model starts with the collection of accurate and sufficient training data. Any biases within the training data can be picked up by the model resulting in inaccurate classification on unseen data. Hence, it is fundamental to understand the biological sequence distribution of the training data and identify any potential biases due to data collection before training and validation. In this paper, our main focus was on the intrinsic diversity of epitope-specific TCR β repertoires. In general, various TCR sequences can recognize the same epitope. These sequences can be either quite similar to each other or contain very distinct features.

When looking at the epitope-specific TCR β repertoire in its entirety, multiple clusters of similar TCRs can be seen spread out over the sequence similarity space. Each of these clusters can be summarized as one motif. In other words, TCRs with different specificities can share similar or identical motifs, while these can be very different for TCRs with the same specificity. This conclusion could also be drawn from the combined clustering of all epitope-specific TCR data within TCRex, where motifs could be shared across clusters with different specificities.

To understand the origin of the clustered TCR space, we simulated epitope-specific TCR repertoires with LIgO. In general, these repertoires were generated by selecting TCRs out of a pool of *in silico* VDJ recombined TCRs that contain a pre-defined motif. In theory, this motif can be considered an epitope-binding motif. Of course, the exact specificity of the simulated TCRs cannot be guaranteed or predicted. However, this strategy allows us to investigate how one epitope-specific binding motif gives rise to a diverse TCR repertoire. Investigation of the simulated repertoires showed that one binding motif can give rise to various TCR clusters, similar to experimental epitope-specific TCR repertoires. This is likely the result of the intrinsic constraints of VDJ recombination underlying the TCR nucleotide sequences. Not all combinations of a motif are possible or highly unlikely due to the finite selection of V and J genes. This produces ‘gaps’ in the generated TCRs, simulated or real, which then lead to their own unique clusters, despite sharing a universal motif. In addition, we showed that negative TCRs can look very similar to epitope-specific TCRs up to only one amino acid difference or identical CDR3 β sequences with different V/J genes. These could further separate the TCRs into clusters. The match between our simulations and experimental epitope-specific TCR repertoires is further evidenced by the similar distribution in simulated TCR clusters derived from different motifs and those presented in the real epitope-specific data sets. Thus, the general sequence distribution of TCR repertoires can be explained by the diversity of epitope-specific motifs within the TCR sequences combined with specific constraints.

The number of clusters and their size can highly differ between epitope-specific TCR repertoires. Here, we showed that the arrangement of the clustered repertoire is a key factor in prediction model performance. In general, clonal TCR repertoires, i.e. having a few large clusters, result in prediction models with better performance than TCR repertoires containing no large clusters. This is in accordance with previous research where a negative correlation between diversity and model performance was demonstrated (4,5,17,18). Different factors might contribute to the variation in the TCR repertoire diversity per epitope. First of all, some epitopes might be more immunodominant than others and thus activate a wider range of T cells (35). In addition, the observed diversity might be affected by the sample collection step. Popular epitopes that are investigated by various research groups will be presented by a substantial collection of TCRs gathered from a larger group of individuals in contrast to the lesser studied epitopes. As a consequence, one might ask if models trained on data collected from a limited number of studies can make reliable predictions for TCR repertoires in new studies. One possible strategy to investigate this is by using a leave-one-study-out (7) or leave-one-subject-out (4,5). in addition to the standard cross-validation. However, it could also be interesting to evaluate the overlap with the training data for each TCR individually by calculating a similarity score with each TCR in the training data or its presence in one of its clusters.

In addition to epitope-specific TCR sequences, training prediction models also require a negative training data set. Here, we demonstrated that this data set might contain TCRs that are very similar to those in the epitope-specific training data set. The higher the overlap in TCR amino acid patterns in the two training datasets, the lower the model performance during cross-validation. Hence, both the sequence distribution of the epitope-specific and the negative training data set are important to consider when training and validating models. Indeed, Montemurro et al. used a similar method to demonstrate a relationship between model performance and the level of discrepancy between the epitope-specific and negative training data (6). This is an intrinsic property of a given epitope, or at least the TCRs that are known to be reactive for it. Some epitopes as shown here have known TCR β chains that match more closely to those commonly found in full non-specific repertoires, and the TCRs for these epitopes are harder to classify irrespective of other parameters. One explanation for the large overlap in epitope-specific and non-specific TCR sequences is the effect of single amino acid changes on the affinity of the TCR for the presented epitope. In case a small variation in the TCR sequence results in a large drop in affinity, these lower-affinity TCRs can be missed during experimental identification, including tetramer-based assays (36,37). Of course, the overlap might also be a result of the presence of false negatives in the negative training data as these TCRs were collected from healthy individuals, which are not exempted from naïve T cells recognizing the epitope. Moreover, the number of false negatives could differ as for some epitopes more specific T cells are found within naïve repertoires than others (14,15). Due to this large overlap in epitope-specific and non-specific TCR sequences, we advise against the removal of similar TCRs in the training data for benchmarking purposes. By reducing the number of positive TCRs in the training data set, it can increase the gap between the positive and negative data resulting in incorrect predictions. In contrast, the use of a similarity metric is recommended to evaluate for each TCR its distance with the closest positive and negative TCR.

Another important point of consideration is the relative importance of the α and β chain in the binding. This can be different per epitope (2). Complexes where the α chains carry out most of the epitope-specific interactions will result in less specific CDR3 β chains entering the training data set. This might additionally influence the model performance.

In conclusion, we showed that a high diversity of epitope-specific TCR β repertoires can be the result of a smaller set of binding motifs. This results in a clustered repertoire composition, which directly influences the epitope-specific prediction performance. The negative training data is located in between these clusters resulting in negative TCRs being either very similar or dissimilar to the epitope-specific training data. This similarity is also related with model performance. Hence the sequence distribution of both epitope-specific and negative data has a strong influence on model performance. It is thus crucial to investigate the composition of the entire training data to understand and correctly interpret model performance.

