## Supplemental Materials for "Revealing the hidden sequence distribution of epitope-specific TCR repertoires and its influence on machine learning model performance"

### S1: Selection of seeds for the simulation of epitope-specific TCR repertoires

Simulating epitope-specific TCR repertoires with LIgO requires a description of the motifs that need to be present in the TCR sequences. Here, a motif was described using a seed, i.e. a short amino acid sequence, and a list of possible hamming distances. The seeds were selected from experimental epitope-specific TCRs present in the VDJdb. Table S1.1 gives an overview of the seeds used in every simulation and the original TCR sequences it was derived from.

**Table S1.1:** Overview of all used LIgO seeds, the original TCR sequences in the VDJdb and additional background information.

| Simulation | Original TCR | LIgO seed | Additional info from VDJdb | | | | |
|---|---|---|---|---|---|---|---|
| | | | V gene | J gene | Epitope | Species | HLA background |
| 1 | CSVWTGEKHEAFF | WTGEKHE | TRBV29-1 | TRBJ1-1 | FLKEKGGL | HIV-1 | HLA-B*08 |
| 1 | CSASSQRGGIYEQYF | SSQRGGIYE | TRBV20-1 | TRBJ2-7 | FLKEKGGL | HIV-1 | HLA-B*08 |
| 1 | CSAHLYRAYGYTF | HLYRAYG | TRBV20-1 | TRBJ1-2 | FLKEKGGL | HIV-1 | HLA-B*08 |
| 2 | CATKGTGLYNEQFF | KGTGLYNE | TRBV24-1 | TRBJ2-1 | FLKEKGGL | HIV-1 | HLA-B*08 |
| 2 | CASSYERGMNTEAFF | SYERGMNTE | TRBV6-3 | TRBJ1-1 | FLKEKGGL | HIV-1 | HLA-B*08 |
| 2 | CASSRAGADYNEQFF | SRAGADYNE | TRBV7-9 | TRBJ2-1 | FLKEKGGL | HIV-1 | HLA-B*08 |
| 3 | CASSLAVGGYEQYF | SLAVGGYE | TRBV11-2 | TRBJ2-7 | GTSGSPIVNR | DENV1 | HLA-A*11 |
| 3 | CSVELSGINQPQHF | ELSGINQP | TRBV29-1 | TRBJ1-5 | GTSGSPIVNR | DENV1 | HLA-A*11 |
| 3 | CASSRGSAETQYF | SRGSAET | TRBV9 | TRBJ2-5 | GTSGSPIVNR | DENV1 | HLA-A*11 |
| 4 | CSAPIPPYNEQFF | PIPPYNE | TRBV20-1 | TRBJ2-1 | YVLDHLIVV | EBV | HLA-A*02 |
| 4 | CSVVLAPVQEQFF | VLAPVQE | TRBV29-1 | TRBJ2-1 | YVLDHLIVV | EBV | HLA-A*02 |
| 4 | CASRTLLGGASEQYF | RTLLGGASE | TRBV27 | TRBJ2-7 | YVLDHLIVV | EBV | HLA-A*02 |
| 5 | CSVPFAGADTQYF | PFAGADT | TRBV29-1 | TRBJ2-3 | LLWNGPMAV | YFV | HLA-A*02 |
| 5 | CSATGNRGADTQYF | TGNRGADT | TRBV20-1 | TRBJ2-3 | LLWNGPMAV | YFV | HLA-A*02 |
| 5 | CASSYSEQGYGYTF | SYSEQGYG | TRBV6-5 | TRBJ1-2 | LLWNGPMAV | YFV | HLA-A*02 |
| 6 | CSATPWGGSYEQYF | TPWGGSYE | TRBV20-1 | TRBJ2-7 | KTFPPTEPK | SARS-CoV-2 | HLA-A*03 |
| 6 | CASSRAAGSKDTQYF | SRAAGSKDT | TRBV4-1 | TRBJ2-3 | KTFPPTEPK | SARS-CoV-2 | HLA-A*03 |
| 6 | CASSHSLAGGSNEQFF | SHSLAGGSNE | TRBV4-3 | TRBJ2-1 | KTFPPTEPK | SARS-CoV-2 | HLA-A*03 |
| 7 | CSGVVGRGAYNEQFF | VVGRGAYNE | TRBV29-1 | TRBJ2-1 | YSEHPTFTSQY | CMV | HLA-A*01 |
| 7 | CASSRKGGGMWTEAFF | SRKGGGMWTE | TRBV12-4 | TRBJ1-1 | YSEHPTFTSQY | CMV | HLA-A*01 |
| 7 | CAISEPTSGRDTQYF | SEPTSGRDT | TRBV10-3 | TRBJ2-3 | YSEHPTFTSQY | CMV | HLA-A*01 |
| 8 | CASSPLLAGGPYEQYF | SPLLAGGPYE | TRBV12-4 | TRBJ2-7 | YSEHPTFTSQY | CMV | HLA-A*01 |
| 8 | CASSLEGDMDSEQYF | SLEGDMDSE | TRBV27 | TRBJ2-7 | YSEHPTFTSQY | CMV | HLA-A*01 |
| 8 | CASNPTGDFYEQYF | NPTGDFYE | TRBV2 | TRBJ2-7 | YSEHPTFTSQY | CMV | HLA-A*01 |

**S2: Overview of all training data and model performances of TCRex epitopes**

TCRex was used to train prediction models for 6 cancer and 120 viral epitopes. The performances of these models are summarized in table S2.1.

**Table S2.1:** Training data size and model performances of TCRex classifiers

| Epitope | Viral/Cancer | Origin | Number of TCR sequences | Balanced accuracy | Average precision | ROC AUC | Active |
|---|---|---|---|---|---|---|---|
| EAAGIGILTV | Cancer | Melanoma | 266 | 0.69 ¬± 0.03 | 0.74 ¬± 0.01 | 0.9 ¬± 0.01 | Yes |
| ELAGIGILTV | Cancer | Melanoma | 1035 | 0.54 ¬± 0.0 | 0.37 ¬± 0.03 | 0.72 ¬± 0.01 | Yes |
| AMFWSVPTV | Cancer | Melanoma | 82 | 0.59 ¬± 0.04 | 0.48 ¬± 0.12 | 0.78 ¬± 0.04 | Yes |
| FLYNLLTRV | Cancer | Melanoma | 61 | 0.63 ¬± 0.08 | 0.59 ¬± 0.1 | 0.87 ¬± 0.03 | Yes |
| LLLGIGILV | Cancer | Multiple Myeloma | 233 | 0.58 ¬± 0.01 | 0.44 ¬± 0.06 | 0.77 ¬± 0.04 | Yes |
| NLSALGIFST | Cancer | Unknown | 111 | 0.51 ¬± 0.01 | 0.19 ¬± 0.03 | 0.66 ¬± 0.04 | No |
| TPRVTGGGAM | Viral | CMV | 258 | 0.7 ¬± 0.02 | 0.75 ¬± 0.04 | 0.88 ¬± 0.03 | Yes |
| NLVPMVATV | Viral | CMV | 4812 | 0.56 ¬± 0.0 | 0.39 ¬± 0.01 | 0.72 ¬± 0.01 | Yes |
| QYDPVAALF | Viral | CMV | 41 | 0.7 ¬± 0.06 | 0.6 ¬± 0.06 | 0.82 ¬± 0.06 | Yes |
| YSEHPTFTSQY | Viral | CMV | 74 | 0.59 ¬± 0.06 | 0.71 ¬± 0.06 | 0.93 ¬± 0.04 | Yes |
| QIKVRVKMV | Viral | CMV | 36 | 0.5 ¬± 0.0 | 0.4 ¬± 0.09 | 0.79 ¬± 0.05 | Yes |
| VTEHDTLLY | Viral | CMV | 277 | 0.54 ¬± 0.0 | 0.39 ¬± 0.04 | 0.78 ¬± 0.02 | Yes |
| IPSINVHHY | Viral | CMV | 85 | 0.7 ¬± 0.02 | 0.66 ¬± 0.06 | 0.84 ¬± 0.04 | Yes |
| MLNIPSINV | Viral | CMV | 73 | 0.51 ¬± 0.01 | 0.25 ¬± 0.04 | 0.61 ¬± 0.04 | No |
| GTSGSPIVNR | Viral | DENV1 | 165 | 0.75 ¬± 0.05 | 0.76 ¬± 0.06 | 0.88 ¬± 0.04 | Yes |
| GTSGSPIIDK | Viral | DENV2 | 60 | 0.61 ¬± 0.06 | 0.49 ¬± 0.04 | 0.74 ¬± 0.08 | Yes |
| GTSGSPIINR | Viral | DENV3/4 | 158 | 0.73 ¬± 0.02 | 0.75 ¬± 0.03 | 0.86 ¬± 0.03 | Yes |
| IVTDFSVIK | Viral | EBV | 46 | 0.6 ¬± 0.05 | 0.53 ¬± 0.05 | 0.83 ¬± 0.04 | Yes |
| RAKFKQLL | Viral | EBV | 262 | 0.65 ¬± 0.02 | 0.69 ¬± 0.04 | 0.89 ¬± 0.01 | Yes |
| GLCTLVAML | Viral | EBV | 1208 | 0.66 ¬± 0.01 | 0.59 ¬± 0.0 | 0.82 ¬± 0.02 | Yes |
| YVLDHLIVV | Viral | EBV | 103 | 0.52 ¬± 0.02 | 0.44 ¬± 0.09 | 0.76 ¬± 0.07 | Yes |
| EPLPQGQLTAY | Viral | EBV | 36 | 0.64 ¬± 0.08 | 0.68 ¬± 0.18 | 0.91 ¬± 0.06 | Yes |
| HPVGEADYFEY | Viral | EBV | 32 | 0.72 ¬± 0.08 | 0.68 ¬± 0.13 | 0.86 ¬± 0.08 | Yes |
| HSKKKCDEL | Viral | HCV | 45 | 0.77 ¬± 0.09 | 0.93 ¬± 0.03 | 0.99 ¬± 0.01 | Yes |
| ATDALMTGY | Viral | HCV | 177 | 0.75 ¬± 0.03 | 0.77 ¬± 0.06 | 0.91 ¬± 0.04 | Yes |
| KLVALGINAV | Viral | HCV | 65 | 0.62 ¬± 0.05 | 0.51 ¬± 0.14 | 0.73 ¬± 0.08 | Yes |
| CINGVCWTV | Viral | HCV | 131 | 0.53 ¬± 0.03 | 0.37 ¬± 0.11 | 0.75 ¬± 0.06 | Yes |
| ARMILMTHF | Viral | HCV | 66 | 0.74 ¬± 0.07 | 0.69 ¬± 0.12 | 0.86 ¬± 0.08 | Yes |
| KAFSPEVIPMF | Viral | HIV | 210 | 0.73 ¬± 0.02 | 0.71 ¬± 0.03 | 0.9 ¬± 0.01 | Yes |
| EIYKRWII | Viral | HIV | 185 | 0.65 ¬± 0.02 | 0.52 ¬± 0.04 | 0.75 ¬± 0.06 | Yes |
| FLKEKGGL | Viral | HIV | 156 | 0.6 ¬± 0.03 | 0.48 ¬± 0.1 | 0.76 ¬± 0.06 | Yes |
| KRWIIMGLNK | Viral | HIV | 75 | 0.69 ¬± 0.05 | 0.73 ¬± 0.12 | 0.85 ¬± 0.09 | Yes |
| ISPRTLNAW | Viral | HIV | 58 | 0.61 ¬± 0.06 | 0.6 ¬± 0.12 | 0.84 ¬± 0.03 | Yes |
| QASQEVKNW | Viral | HIV | 31 | 0.58 ¬± 0.05 | 0.31 ¬± 0.12 | 0.6 ¬± 0.11 | No |
| TPQDLNTML | Viral | HIV | 159 | 0.81 ¬± 0.02 | 0.87 ¬± 0.06 | 0.94 ¬± 0.04 | Yes |
| KRWIILGLNK | Viral | HIV | 396 | 0.65 ¬± 0.02 | 0.64 ¬± 0.08 | 0.85 ¬± 0.04 | Yes |
| RYPLTFGWCF | Viral | HIV | 30 | 0.52 ¬± 0.03 | 0.42 ¬± 0.12 | 0.72 ¬± 0.13 | Yes |
| FPRPWLHGL | Viral | HIV | 120 | 0.81 ¬± 0.02 | 0.83 ¬± 0.03 | 0.93 ¬± 0.02 | Yes |
| IIKDYGKQM | Viral | HIV | 54 | 0.81 ¬± 0.07 | 0.84 ¬± 0.07 | 0.95 ¬± 0.04 | Yes |
| LPPIVAKEI | Viral | HIV | 62 | 0.79 ¬± 0.04 | 0.78 ¬± 0.06 | 0.9 ¬± 0.05 | Yes |
| HPKVSSEVHI | Viral | HIV | 75 | 0.69 ¬± 0.07 | 0.7 ¬± 0.16 | 0.87 ¬± 0.08 | Yes |
| TPGPGVRYPL | Viral | HIV | 86 | 0.68 ¬± 0.04 | 0.74 ¬± 0.07 | 0.88 ¬± 0.03 | Yes |
| RFYKTLRAEQASQ | Viral | HIV | 210 | 0.9 ¬± 0.03 | 0.96 ¬± 0.01 | 0.99 ¬± 0.0 | Yes |
| SLYNTVATL | Viral | HIV | 58 | 0.63 ¬± 0.03 | 0.4 ¬± 0.05 | 0.59 ¬± 0.06 | No |
| GPGHKARVL | Viral | HIV | 66 | 0.51 ¬± 0.02 | 0.31 ¬± 0.11 | 0.67 ¬± 0.06 | No |
| RLRPGGKKK | Viral | HIV | 31 | 0.73 ¬± 0.06 | 0.73 ¬± 0.2 | 0.89 ¬± 0.09 | Yes |
| QVPLRPMTYK | Viral | HIV | 48 | 0.64 ¬± 0.04 | 0.51 ¬± 0.16 | 0.77 ¬± 0.11 | Yes |
| FRDYVDRFYKTLRAEQASQE | Viral | HIV | 367 | 0.87 ¬± 0.02 | 0.96 ¬± 0.01 | 1.0 ¬± 0.0 | Yes |
| RPRGEVRFL | Viral | HSV2 | 63 | 0.77 ¬± 0.04 | 0.84 ¬± 0.08 | 0.92 ¬± 0.05 | Yes |
| SFHSLHLLF | Viral | HTLV1 | 131 | 0.68 ¬± 0.03 | 0.63 ¬± 0.08 | 0.81 ¬± 0.08 | Yes |
| PKYVKQNTLKLAT | Viral | Influenza | 292 | 0.52 ¬± 0.01 | 0.36 ¬± 0.05 | 0.72 ¬± 0.02 | Yes |
| GILGFVFTL | Viral | Influenza | 3107 | 0.68 ¬± 0.01 | 0.59 ¬± 0.02 | 0.81 ¬± 0.02 | Yes |
| LPRRSGAAGA | Viral | Influenza | 2137 | 0.54 ¬± 0.01 | 0.4 ¬± 0.02 | 0.84 ¬± 0.01 | Yes |
| FLNRFTTTL | Viral | SARS-CoV-2 | 104 | 0.54 ¬± 0.03 | 0.38 ¬± 0.12 | 0.72 ¬± 0.07 | Yes |
| GTHWFVTQR | Viral | SARS-CoV-2 | 98 | 0.5 ¬± 0.0 | 0.19 ¬± 0.04 | 0.66 ¬± 0.04 | No |
| TTLPVNVAF | Viral | SARS-CoV-2 | 31 | 0.52 ¬± 0.03 | 0.26 ¬± 0.07 | 0.65 ¬± 0.06 | No |
| NYSGVVTTVMF | Viral | SARS-CoV-2 | 33 | 0.51 ¬± 0.03 | 0.29 ¬± 0.05 | 0.68 ¬± 0.12 | No |
| HTDFSSEIIGY | Viral | SARS-CoV-2 | 35 | 0.6 ¬± 0.06 | 0.54 ¬± 0.1 | 0.67 ¬± 0.09 | No |
| WICLLQFAY | Viral | SARS-CoV-2 | 515 | 0.64 ¬± 0.02 | 0.56 ¬± 0.06 | 0.81 ¬± 0.02 | Yes |
| HLVDFQVTI | Viral | SARS-CoV-2 | 69 | 0.61 ¬± 0.04 | 0.44 ¬± 0.09 | 0.75 ¬± 0.06 | Yes |
| KLNVGDYFV | Viral | SARS-CoV-2 | 143 | 0.52 ¬± 0.02 | 0.31 ¬± 0.07 | 0.72 ¬± 0.04 | No |

| Epitope | Viral/Cancer | Origin | Number of TCR sequences | Balanced accuracy | Average precision | ROC AUC | Active |
|---|---|---|---|---|---|---|---|
| TFYLTNDVSFL | Viral | SARS-CoV-2 | 33 | 0.57 ¬± 0.08 | 0.43 ¬± 0.1 | 0.66 ¬± 0.04 | No |
| LPAADLDDF | Viral | SARS-CoV-2 | 131 | 0.53 ¬± 0.02 | 0.32 ¬± 0.04 | 0.73 ¬± 0.02 | No |
| EILDITPCSF | Viral | SARS-CoV-2 | 68 | 0.63 ¬± 0.03 | 0.46 ¬± 0.09 | 0.73 ¬± 0.06 | Yes |
| YFPLQSYGF | Viral | SARS-CoV-2 | 357 | 0.53 ¬± 0.01 | 0.4 ¬± 0.04 | 0.79 ¬± 0.03 | Yes |
| AYILFTRFFYV | Viral | SARS-CoV-2 | 119 | 0.52 ¬± 0.01 | 0.28 ¬± 0.07 | 0.7 ¬± 0.05 | No |
| LVLSVNPYV | Viral | SARS-CoV-2 | 50 | 0.5 ¬± 0.0 | 0.34 ¬± 0.09 | 0.66 ¬± 0.11 | No |
| FTISVTTEIL | Viral | SARS-CoV-2 | 159 | 0.55 ¬± 0.03 | 0.37 ¬± 0.04 | 0.73 ¬± 0.03 | Yes |
| LPPAYTNSF | Viral | SARS-CoV-2 | 127 | 0.53 ¬± 0.02 | 0.45 ¬± 0.06 | 0.82 ¬± 0.03 | Yes |
| QECVRGTTVL | Viral | SARS-CoV-2 | 147 | 0.71 ¬± 0.02 | 0.66 ¬± 0.04 | 0.85 ¬± 0.02 | Yes |
| MPASWVMRI | Viral | SARS-CoV-2 | 477 | 0.62 ¬± 0.02 | 0.53 ¬± 0.01 | 0.82 ¬± 0.01 | Yes |
| VLWAHGFEL | Viral | SARS-CoV-2 | 695 | 0.65 ¬± 0.02 | 0.6 ¬± 0.04 | 0.85 ¬± 0.02 | Yes |
| KEIDRLNEV | Viral | SARS-CoV-2 | 57 | 0.58 ¬± 0.02 | 0.43 ¬± 0.1 | 0.7 ¬± 0.08 | Yes |
| TPINLVRDL | Viral | SARS-CoV-2 | 249 | 0.64 ¬± 0.02 | 0.53 ¬± 0.04 | 0.81 ¬± 0.03 | Yes |
| KPLEFGATSAAL | Viral | SARS-CoV-2 | 344 | 0.59 ¬± 0.02 | 0.49 ¬± 0.07 | 0.8 ¬± 0.04 | Yes |
| VLAWLYAAV | Viral | SARS-CoV-2 | 112 | 0.5 ¬± 0.01 | 0.18 ¬± 0.05 | 0.6 ¬± 0.04 | No |
| FIAGLIAIV | Viral | SARS-CoV-2 | 184 | 0.52 ¬± 0.02 | 0.29 ¬± 0.08 | 0.71 ¬± 0.04 | No |
| YIFFASFYY | Viral | SARS-CoV-2 | 272 | 0.5 ¬± 0.0 | 0.31 ¬± 0.03 | 0.77 ¬± 0.02 | No |
| RQLLFVVEV | Viral | SARS-CoV-2 | 841 | 0.57 ¬± 0.01 | 0.46 ¬± 0.02 | 0.83 ¬± 0.01 | Yes |
| ALSKGVHFV | Viral | SARS-CoV-2 | 129 | 0.53 ¬± 0.01 | 0.4 ¬± 0.09 | 0.79 ¬± 0.03 | Yes |
| FLPRVFSAV | Viral | SARS-CoV-2 | 773 | 0.59 ¬± 0.02 | 0.51 ¬± 0.03 | 0.84 ¬± 0.01 | Yes |
| SLVKPSFYV | Viral | SARS-CoV-2 | 50 | 0.52 ¬± 0.02 | 0.38 ¬± 0.08 | 0.69 ¬± 0.03 | No |
| IPRRNVATL | Viral | SARS-CoV-2 | 48 | 0.57 ¬± 0.02 | 0.32 ¬± 0.08 | 0.58 ¬± 0.09 | No |
| IQYIDIGNY | Viral | SARS-CoV-2 | 149 | 0.56 ¬± 0.03 | 0.47 ¬± 0.06 | 0.82 ¬± 0.04 | Yes |
| KLSYGIATV | Viral | SARS-CoV-2 | 2149 | 0.58 ¬± 0.0 | 0.51 ¬± 0.01 | 0.84 ¬± 0.01 | Yes |
| ILHCANFNV | Viral | SARS-CoV-2 | 185 | 0.61 ¬± 0.02 | 0.52 ¬± 0.05 | 0.84 ¬± 0.03 | Yes |
| KLWAQCVQL | Viral | SARS-CoV-2 | 266 | 0.55 ¬± 0.01 | 0.46 ¬± 0.05 | 0.8 ¬± 0.02 | Yes |
| SEISMDNSPNL | Viral | SARS-CoV-2 | 95 | 0.52 ¬± 0.01 | 0.45 ¬± 0.13 | 0.79 ¬± 0.05 | Yes |
| KTSVDCTMYI | Viral | SARS-CoV-2 | 70 | 0.7 ¬± 0.05 | 0.72 ¬± 0.08 | 0.89 ¬± 0.05 | Yes |
| IVDTVSALV | Viral | SARS-CoV-2 | 36 | 0.51 ¬± 0.03 | 0.36 ¬± 0.18 | 0.76 ¬± 0.12 | Yes |
| LLFNKVTLA | Viral | SARS-CoV-2 | 39 | 0.51 ¬± 0.03 | 0.37 ¬± 0.1 | 0.76 ¬± 0.04 | Yes |
| SEETGTLIV | Viral | SARS-CoV-2 | 38 | 0.6 ¬± 0.03 | 0.39 ¬± 0.08 | 0.68 ¬± 0.09 | No |
| TLDSKTQSL | Viral | SARS-CoV-2 | 107 | 0.83 ¬± 0.06 | 0.88 ¬± 0.06 | 0.97 ¬± 0.02 | Yes |
| TVYDPLQPELDSFK | Viral | SARS-CoV-2 | 36 | 0.51 ¬± 0.03 | 0.23 ¬± 0.08 | 0.53 ¬± 0.07 | No |
| FPPTSFGPL | Viral | SARS-CoV-2 | 621 | 0.69 ¬± 0.01 | 0.61 ¬± 0.01 | 0.85 ¬± 0.01 | Yes |
| SSNVANYQK | Viral | SARS-CoV-2 | 74 | 0.61 ¬± 0.04 | 0.45 ¬± 0.08 | 0.77 ¬± 0.04 | Yes |
| YLNTLTLAV | Viral | SARS-CoV-2 | 390 | 0.58 ¬± 0.02 | 0.46 ¬± 0.04 | 0.8 ¬± 0.01 | Yes |
| LLMPILTLT | Viral | SARS-CoV-2 | 46 | 0.5 ¬± 0.0 | 0.25 ¬± 0.06 | 0.69 ¬± 0.06 | No |
| NQKLIANQF | Viral | SARS-CoV-2 | 51 | 0.76 ¬± 0.02 | 0.78 ¬± 0.07 | 0.95 ¬± 0.02 | Yes |
| ILGLPTQTV | Viral | SARS-CoV-2 | 198 | 0.68 ¬± 0.02 | 0.6 ¬± 0.08 | 0.83 ¬± 0.05 | Yes |
| YLQPRTFLL | Viral | SARS-CoV-2 | 315 | 0.91 ¬± 0.01 | 0.92 ¬± 0.01 | 0.97 ¬± 0.01 | Yes |
| YLDAYNMMI | Viral | SARS-CoV-2 | 197 | 0.62 ¬± 0.03 | 0.42 ¬± 0.05 | 0.74 ¬± 0.03 | Yes |
| SEPVLKGVKL | Viral | SARS-CoV-2 | 80 | 0.57 ¬± 0.02 | 0.38 ¬± 0.1 | 0.79 ¬± 0.05 | Yes |
| TLIGDCATV | Viral | SARS-CoV-2 | 467 | 0.55 ¬± 0.02 | 0.36 ¬± 0.04 | 0.73 ¬± 0.03 | Yes |
| ITEEVGHTDLMAAY | Viral | SARS-CoV-2 | 156 | 0.57 ¬± 0.02 | 0.49 ¬± 0.04 | 0.78 ¬± 0.02 | Yes |
| ISDYDYYRY | Viral | SARS-CoV-2 | 39 | 0.5 ¬± 0.0 | 0.25 ¬± 0.09 | 0.67 ¬± 0.06 | No |
| EEHVQIHTI | Viral | SARS-CoV-2 | 104 | 0.5 ¬± 0.01 | 0.2 ¬± 0.09 | 0.59 ¬± 0.07 | No |
| IPIQASLPF | Viral | SARS-CoV-2 | 104 | 0.51 ¬± 0.02 | 0.35 ¬± 0.11 | 0.74 ¬± 0.09 | Yes |
| TLVPQEHYV | Viral | SARS-CoV-2 | 154 | 0.53 ¬± 0.03 | 0.42 ¬± 0.07 | 0.72 ¬± 0.02 | Yes |
| FADDLNQLTGY | Viral | SARS-CoV-2 | 70 | 0.61 ¬± 0.04 | 0.44 ¬± 0.11 | 0.7 ¬± 0.05 | Yes |
| RIFTIGTVTLK | Viral | SARS-CoV-2 | 81 | 0.51 ¬± 0.02 | 0.32 ¬± 0.15 | 0.67 ¬± 0.1 | No |
| YEGNSPFHPL | Viral | SARS-CoV-2 | 60 | 0.53 ¬± 0.03 | 0.31 ¬± 0.06 | 0.66 ¬± 0.09 | No |
| NLNESLIDL | Viral | SARS-CoV-2 | 132 | 0.59 ¬± 0.03 | 0.42 ¬± 0.05 | 0.74 ¬± 0.02 | Yes |
| SEVGPEHSLAEY | Viral | SARS-CoV-2 | 250 | 0.54 ¬± 0.01 | 0.42 ¬± 0.05 | 0.79 ¬± 0.01 | Yes |
| NLDSKVGGNY | Viral | SARS-CoV-2 | 45 | 0.73 ¬± 0.06 | 0.85 ¬± 0.05 | 0.96 ¬± 0.03 | Yes |
| TSNQVAVLY | Viral | SARS-CoV-2 | 62 | 0.52 ¬± 0.02 | 0.25 ¬± 0.07 | 0.69 ¬± 0.09 | No |
| KAYNVTQAF | Viral | SARS-CoV-2 | 708 | 0.61 ¬± 0.02 | 0.57 ¬± 0.03 | 0.83 ¬± 0.01 | Yes |
| GVAMPNLYK | Viral | SARS-CoV-2 | 35 | 0.5 ¬± 0.0 | 0.12 ¬± 0.02 | 0.54 ¬± 0.1 | No |
| LEPLVDLPI | Viral | SARS-CoV-2 | 367 | 0.54 ¬± 0.01 | 0.35 ¬± 0.05 | 0.76 ¬± 0.02 | Yes |
| KLPDDFTGCV | Viral | SARS-CoV-2 | 1160 | 0.59 ¬± 0.01 | 0.54 ¬± 0.02 | 0.84 ¬± 0.01 | Yes |
| FLNGSCGSV | Viral | SARS-CoV-2 | 2332 | 0.61 ¬± 0.0 | 0.56 ¬± 0.02 | 0.86 ¬± 0.01 | Yes |
| FVDGVPFVV | Viral | SARS-CoV-2 | 2420 | 0.54 ¬± 0.0 | 0.37 ¬± 0.01 | 0.77 ¬± 0.01 | Yes |
| HTTDPSFLGRY | Viral | SARS-CoV-2 | 5000 | 0.7 ¬± 0.01 | 0.7 ¬± 0.02 | 0.9 ¬± 0.01 | Yes |
| ILIEGIFFV | Viral | VZV | 111 | 0.63 ¬± 0.03 | 0.55 ¬± 0.09 | 0.82 ¬± 0.04 | Yes |
| ALSQYHVYV | Viral | VZV | 69 | 0.62 ¬± 0.03 | 0.51 ¬± 0.06 | 0.74 ¬± 0.04 | Yes |
| LLWNGPMAV | Viral | YellowFeverVirus | 474 | 0.66 ¬± 0.01 | 0.53 ¬± 0.03 | 0.79 ¬± 0.01 | Yes |

Note that for one epitope (i.e. NLSALGIFST) the exact cancer pathology was not available in the original public VDJdb database and is thus associated with an 'Unknown' origin.

**S3: Clustering of TCRex training data**

All TCRex training data was clustered together with ClusTCR. This resulted in clusters of various sizes as shown in table S3.1. A subset of these were pure, i.e. they contained CDR3 β sequences associated with the same epitope. The number and sizes of these pure clusters are listed in the file results/tcrex_clustering/pure_clusters.tsv.

**Table S3.1:** Overview of the sizes of all clusters found in the TCRex training data

| Size | Count |
|---|---|
| 2 | 1298 |
| 3 | 335 |
| 4 | 134 |
| 5 | 59 |
| ]5, 10] | 109 |
| ]10, 15] | 40 |
| ]15, 20] | 28 |
| ]20, 50] | 65 |
| ]50, 100] | 27 |
| ]100, 200] | 22 |
| ]200, 300] | 2 |
| ]300, 400] | 2 |

**S4: Clustering of epitope-specific TCR motifs**

Clustering each epitope-specific TCR repertoire in TCRex resulted in a list of epitope-specific motifs. In the following step, all motifs were clustered using the same strategy. The results are stored in 'results/epitope_specific_clustering/cluster_motifs/motif_clusters.tsv'. This file contains for each cluster, the motifs and their associated epitopes.

The file 'results/epitope_specific_clustering/overlapping_motifs.tsv' lists those motifs that were present in different epitope-specific TCR repertoires

**S5: Overlap between epitope-specific and negative TCRs**

For each epitope, the positive and negative training data were clustered:
- The file results/background/shared_clusters.tsv lists for each epitope how many of the clusters contain both epitope-specific and negative TCRs.

- The file results/background/shared_motifs.tsv lists overlapping motifs of epitope-specific and negative clusters. Here the epitope-specific and negative training data was thus clustered separately

In addition, the minimal distance between the two training data sets was calculated:
- The file results/background/min_distances.tsv gives an overview of the minimal distance between the background TCRs and the epitope-specific TCRs for every TCRex model. Also the number of TCRs having this minimal distance are reported.

**S6: Clustering of LIgO simulated repertoires**

Table S6.1 gives an overview of the number of TCRs that were clustered in each of the eight simulated repertoires. Table S6.2 gives a more detailed overview of the motifs of these clusters. Each row lists the characteristics of one motif: the consensus motif itself, the LIgO seeds that were used to simulate the TCRs within the cluster and the size of the cluster the consensus motif is derived from.

**Table S6.1:** Clustering statistics for the eight simulated TCR repertoires, where each repertoire is derived from three seeds extracted from the TCRs of a single epitope. For each simulated repertoire, the number of unique CDR3 β sequences and the number of clustered sequences is given. These were used to calculate the percentage of clustered CDR3 β sequences. In addition, the number of clusters is listed.

| Simulation | Number of clustered CDR3 β sequences | Data size | Number of clusters | Percentage of clustered CDR3 β sequences |
|---|---|---|---|---|
| simulation1 | 175 | 862 | 57 | 20.3 |
| simulation2 | 209 | 890 | 53 | 23.5 |
| simulation3 | 237 | 852 | 48 | 27.8 |
| simulation4 | 110 | 883 | 34 | 12.5 |
| simulation5 | 197 | 852 | 41 | 23.1 |
| simulation6 | 227 | 861 | 58 | 26.4 |
| simulation7 | 180 | 887 | 38 | 20.3 |
| simulation8 | 181 | 885 | 53 | 20.5 |

**Table S6.2:** Overview of the 378 unique consensus motifs identified after clustering the eight simulated TCR repertoires separately. The blue rows highlight the clusters containing CDR3 β sequences derived from more than one LIgO seed within one simulation, while the yellow rows highlight the clusters that were derived during different simulations, but share the same consensus motif. This can be explained by the presence of the same CDR3 β sequences within their clusters (table S6.3)

| **Consensus motif** | **LIgO seed** | **Cluster size** | **Simulation** |
|---|---|---|---|
| CAGXRGADTQYF | TGNRGADT | 2 | simulation5 |
| CAPTGXFYEQYF | NPTGDFYE | 2 | simulation8 |
| CARGLAGGAXETQYF | RTLLGGASE | 2 | simulation4 |
| CARPTGXFYEQYF | NPTGDFYE | 2 | simulation8 |
| CASGXRGADTQYF | TGNRGADT | 2 | simulation5 |
| CASIPXYNEQFF | PIPPYNE | 2 | simulation4 |
| CASLXGGGYEQYF | SLAVGGYE | 2 | simulation3 |
| CASLXGINQPQHF | ELSGINQP | 2 | simulation3 |
| CASLXRAYGYTF | HLYRAYG | 2 | simulation1 |
| CASNPTGRXYEQYF | NPTGDFYE | 4 | simulation8 |
| CASNPTGXAYEQYF | NPTGDFYE | 2 | simulation8 |
| CASPGPXYNEQFF | PIPPYNE | 2 | simulation4 |
| CASPPXGGADTQYF | PFAGADT | 2 | simulation5 |
| CASPTGXFYEQYF | NPTGDFYE | 3 | simulation8 |
| CASPXGGSYEQYF | TPWGGSYE | 2 | simulation6 |

| | | | |
|---|---|---|---|
| CASRAAXXTDTQYF | SRAAGSKDT | 3 | simulation6 |
| CASRAGAXLNTEAFF | SRAGADYNE | 2 | simulation2 |
| CASRAGGXTDTQYF | SRAAGSKDT | 4 | simulation6 |
| CASRAGXSTDTQYF | SRAAGSKDT | 3 | simulation6 |
| CASRASGXYNEQFF | SRAGADYNE | 2 | simulation2 |
| CASRDRGXNTEAFF | SYERGMNTE | 2 | simulation2 |
| CASRELAGXNQPQHF | ELSGINQP | 2 | simulation3 |
| CASRGGXXYNEQFF | SRAGADYNE | 3 | simulation2 |
| CASRGVLAGXQETQYF | VLAPVQE | 2 | simulation4 |
| CASRGXNEKLFF | SRGSAET | 2 | simulation3 |
| CASRGXYEQYF | SRGSAET | 2 | simulation3 |
| CASRKGGGXTGELFF | SRKGGGMWTE | 2 | simulation7 |
| CASRKGQGXNTEAFF | SRKGGGMWTE | 2 | simulation7 |
| CASRKXGGGNTEAFF | SRKGGGMWTE | 2 | simulation7 |
| CASRLAGXSDTQYF | SRAAGSKDT | 2 | simulation6 |
| CASRNPTGXSYEQYF | NPTGDFYE | 2 | simulation8 |
| CASRRLXGGAYEQYF | RTLLGGASE | 2 | simulation4 |
| CASRSGXAYNEQFF | SRAGADYNE | 2 | simulation2 |
| CASRSRGGXYEQYF | SSQRGGIYE | 2 | simulation1 |
| CASRTGXSYNEQFF | SRAGADYNE | 2 | simulation2 |
| CASRTLAGGXNEQFF | RTLLGGASE | 2 | simulation4 |
| CASRTXGSTDTQYF | SRAAGSKDT | 2 | simulation6 |
| CASRXGGGGDTEAFF | SRKGGGMWTE | 2 | simulation7 |
| CASRXGGGLNTEAFF | SRKGGGMWTE | 2 | simulation7 |
| CASRXGGGRATEAFF | SRKGGGMWTE | 2 | simulation7 |
| CASRXGGGRNTEAFF | SRKGGGMWTE | 4 | simulation7 |
| CASRXGGPYNEQFF | SRAGADYNE | 4 | simulation2 |
| CASRXGGXMNTEAFF | SRKGGGMWTE | 16 | simulation7 |
| CASRXGTGXYNEQFF | KGTGLYNE | 3 | simulation2 |
| CASRXGTSYNEQFF | SRAGADYNE | 2 | simulation2 |
| CASRXGXGMNTEAFF | SRKGGGMWTE | 5, 10 | simulation7 |
| CASRXLAGGXNEQFF | SHSLAGGSNE | 3 | simulation6 |
| CASRXLGGMNTEAFF | SRKGGGMWTE | 2 | simulation7 |
| CASRXRATYNEQFF | SRAGADYNE | 2 | simulation2 |
| CASRXSYEQYF | SRGSAET | 2 | simulation3 |
| CASRXXGGMNTEAFF | SRKGGGMWTE | 10 | simulation7 |
| CASRXXGSTDTQYF | SRAAGSKDT | 11 | simulation6 |
| CASSAAXSTDTQYF | SRAAGSKDT | 2 | simulation6 |
| CASSAXGSTDTQYF | SRAAGSKDT | 2 | simulation6 |
| CASSAXLAGGPHEQFF | SPLLAGGPYE | 2 | simulation8 |
| CASSCRGGXHEQFF | SSQRGGIYE | 2 | simulation1 |
| CASSDPXRGADTQYF | PFAGADT | 2 | simulation5 |

| | | | |
|---|---|---|---|
| CASSDXXGMNTEAFF | SYERGMNTE | 3 | simulation2 |
| CASSELXGENQPQHF | ELSGINQP | 2 | simulation3 |
| CASSEXSGSNQPQHF | ELSGINQP | 5 | simulation3 |
| CASSEXSGXNQPQHF | ELSGINQP | 3 | simulation3 |
| CASSEXTGINQPQHF | ELSGINQP | 2 | simulation3 |
| CASSFRGAXYNEQFF | SRAGADYNE | 2 | simulation2 |
| CASSFRGXPYEQYF | SSQRGGIYE | 2 | simulation1 |
| CASSFXGADTQYF | PFAGADT | 3 | simulation5 |
| CASSFXRXMNTEAFF | SYERGMNTE | 3 | simulation2 |
| CASSFXTGDFYEQYF | NPTGDFYE | 2 | simulation8 |
| CASSGKGTGXYNSPLHF | KGTGLYNE | 2 | simulation2 |
| CASSGREGXYEQYF | SSQRGGIYE | 2 | simulation1 |
| CASSGRXGRYEQYF | SSQRGGIYE | 2 | simulation1 |
| CASSGTGXRGTDTQYF | TGNRGADT | 2 | simulation5 |
| CASSHLXRAYEQYF | HLYRAYG | 2 | simulation1 |
| CASSHLXRNYGYTF | HLYRAYG | 2 | simulation1 |
| CASSHPLAGGXYEQYF | SHSLAGGSNE | 2 | simulation6 |
| CASSHXSRAYGYTF | HLYRAYG | 2 | simulation1 |
| CASSLAGGXNEKLFF | SHSLAGGSNE | 2 | simulation6 |
| CASSLAGGXNEQFF | SHSLAGGSNE | 3 | simulation6 |
| CASSLAGXKDTQYF | SRAAGSKDT | 2 | simulation6 |
| CASSLAGXXYNEQFF | SRAGADYNE | 3 | simulation2 |
| CASSLARXGYEQFF | SLAVGGYE | 2 | simulation3 |
| CASSLAXGGHEQYF | SLAVGGYE | 2 | simulation3 |
| CASSLAXGGNEKLFF | SLAVGGYE | 2 | simulation3 |
| CASSLAXGGPYEQYF | SPLLAGGPYE | 3 | simulation8 |
| CASSLAXGGQETQYF | SLAVGGYE | 2 | simulation3 |
| CASSLAXGGTEAFF | SLAVGGYE | 3 | simulation3 |
| CASSLAXGGYGYTF | SLAVGGYE | 3 | simulation3 |
| CASSLAXGGYNEQFF | SLAVGGYE | 2 | simulation3 |
| CASSLAXGGYTF | SLAVGGYE | 3 | simulation3 |
| CASSLAXXGYEQYF | SLAVGGYE | 20 | simulation3 |
| CASSLEGDGXSYEQYF | SLEGDMDSE | 2 | simulation8 |
| CASSLEGDRGXETQYF | SLEGDMDSE | 2 | simulation8 |
| CASSLEGDXDTQYF | SLEGDMDSE | 2 | simulation8 |
| CASSLEGDXSSYEQYF | SLEGDMDSE | 2 | simulation8 |
| CASSLEGDXTGELFF | SLEGDMDSE | 2 | simulation8 |
| CASSLEGDXXTEAFF | SLEGDMDSE | 4 | simulation8 |
| CASSLEGDXXYEQYF | SLEGDMDSE | 11 | simulation8 |
| CASSLEGDXYNEQFF | SLEGDMDSE | 2 | simulation8 |
| CASSLEGGXDTEAFF | SLEGDMDSE | 2 | simulation8 |
| CASSLEGGXDYEQXF | SLEGDMDSE | 3 | simulation8 |

| | | | |
|---|---|---|---|
| CASSLEGRXDTEAFF | SLEGDMDSE | 2 | simulation8 |
| CASSLEGXMNTEAFF | SLEGDMDSE | 7 | simulation8 |
| CASSLEGXRDTEAFF | SLEGDMDSE | 2 | simulation8 |
| CASSLELXGXNQPQHF | ELSGINQP | 3 | simulation3 |
| CASSLFWTGEXYEQYF | WTGEKHE | 2 | simulation1 |
| CASSLGAGXPYEQYF | SPLLAGGPYE | 2 | simulation8 |
| CASSLGGASEQXF | RTLLGGASE | 2 | simulation4 |
| CASSLGGAXYNEQFF | SRAGADYNE | 2 | simulation2 |
| CASSLGGXGLYNEQFF | KGTGLYNE | 2 | simulation2 |
| CASSLGPXGGSYEQYF | TPWGGSYE | 2 | simulation6 |
| CASSLGPXRGAYNEQFF | VVGRGAYNE | 2 | simulation7 |
| CASSLGTSGXDTQYF | SEPTSGRDT | 2 | simulation7 |
| CASSLLAGGXYEQYF | SPLLAGGPYE | 9 | simulation8 |
| CASSLLAGPXYEQYF | SPLLAGGPYE | 2 | simulation8 |
| CASSLLAGXPXEQFF | SPLLAGGPYE | 3 | simulation8 |
| CASSLLAGXXYEQYF | SPLLAGGPYE | 3 | simulation8 |
| CASSLLGGXSETQYF | RTLLGGASE | 2 | simulation4 |
| CASSLRGXXYEQYF | SSQRGGIYE | 3 | simulation1 |
| CASSLRPGXYEQYF | SSQRGGIYE | 2 | simulation1 |
| CASSLSGXNQPQHF | ELSGINQP | 16 | simulation3 |
| CASSLTGGXGADTQYF | TGNRGADT | 2 | simulation5 |
| CASSLTXGGPYEQYF | SPLLAGGPYE | 2 | simulation8 |
| CASSLVGGXYEQYF | SSQRGGIYE | 2 | simulation1 |
| CASSLVGXGAYNEQFF | VVGRGAYNE | 2 | simulation7 |
| CASSLWTGEXTEAFF | WTGEKHE | 2 | simulation1 |
| CASSLXGDMNTEAFF | SLEGDMDSE | 4 | simulation8 |
| CASSLXGDRDSNQPQHF | SLEGDMDSE | 2 | simulation8 |
| CASSLXGDRDSPLHF | SLEGDMDSE | 2 | simulation8 |
| CASSLXGDRDTEAFF | SLEGDMDSE | 7 | simulation8 |
| CASSLXGGRGADTQYF | TGNRGADT | 2 | simulation5 |
| CASSLXGGSNEKLFF | SHSLAGGSNE | 2 | simulation6 |
| CASSLXGGSNEQFF | SHSLAGGSNE | 2 | simulation6 |
| CASSLXGGXYEQYF | SSQRGGIYE | 8 | simulation1 |
| CASSLXGINQPQHF | ELSGINQP | 9 | simulation3 |
| CASSLXLAGGXNEQFF | SHSLAGGSNE | 4 | simulation6 |
| CASSLXLYRGYGYTF | HLYRAYG | 2 | simulation1 |
| CASSLXPYRAYGYTF | HLYRAYG | 2 | simulation1 |
| CASSLXRGXNTEAFF | SYERGMNTE | 11 | simulation2 |
| CASSLXVGGKETQYF | SLAVGGYE | 2 | simulation3 |
| CASSLXVLAPYNEQFF | VLAPVQE | 2 | simulation4 |
| CASSLXXGMNTEAFF | SYERGMNTE | 5 | simulation2 |
| CASSLXXGXYEQYF | SRGSAET,SLAVGGYE | 80 | simulation3 |

| | | | |
|---|---|---|---|
| CASSLYRXYGYTF | HLYRAYG | 4 | simulation1 |
| CASSLYXAYGYTF | HLYRAYG | 2 | simulation1 |
| CASSNPTGXXYEQYF | NPTGDFYE | 3 | simulation8 |
| CASSPGLAGGXNEQFF | SPLLAGGPYE,SHSLAGGSNE | 4, 3 | simulation6,simulation8 |
| CASSPGLAGGXYEQYF | SPLLAGGPYE | 2 | simulation8 |
| CASSPGPXYNEQFF | PIPPYNE | 2 | simulation4 |
| CASSPLAGADTGELFF | PFAGADT | 4 | simulation5 |
| CASSPPLAGXPYEQYF | SPLLAGGPYE | 2 | simulation8 |
| CASSPRGGXTEAFF | SSQRGGIYE | 2 | simulation1 |
| CASSPSXPYNEQFF | PIPPYNE | 2 | simulation4 |
| CASSPTSGRXYEQYF | SEPTSGRDT | 2 | simulation7 |
| CASSPWGXSYEQYF | TPWGGSYE | 2 | simulation6 |
| CASSPXGDFYEQYF | NPTGDFYE | 3 | simulation8 |
| CASSPXGGSYEQYF | TPWGGSYE,SHSLAGGSNE | 19 | simulation6 |
| CASSQDLXRAYGYTF | HLYRAYG | 2 | simulation1 |
| CASSQDRGXNTEAFF | SYERGMNTE | 3 | simulation2 |
| CASSQDRGXYEQYF | SSQRGGIYE | 5 | simulation1 |
| CASSQELGGXNQPQHF | ELSGINQP | 2 | simulation3 |
| CASSQELXGXNQPQHF | ELSGINQP | 3 | simulation3 |
| CASSQEXSGXNQPQHF | ELSGINQP | 3 | simulation3 |
| CASSQGGXSYEQYF | SSQRGGIYE | 2 | simulation1 |
| CASSQGLSGXNQPQHF | ELSGINQP | 2 | simulation3 |
| CASSQGXDYNEQFF | SRAGADYNE | 4 | simulation2 |
| CASSQXGGAYEQYF | SSQRGGIYE | 2 | simulation1 |
| CASSQXGQGYEQYF | SSQRGGIYE | 2 | simulation1 |
| CASSQXGXCYEQYF | SSQRGGIYE | 3 | simulation1 |
| CASSQXLAGADTQYF | PFAGADT | 2 | simulation5 |
| CASSQXPLXGINQPQHF | ELSGINQP | 3 | simulation3 |
| CASSRAXQXYNEQFF | SRAGADYNE | 3 | simulation2 |
| CASSRGGPXYNEQFF | SRAGADYNE | 2 | simulation2 |
| CASSRGGXXYNEQFF | SRAGADYNE | 3 | simulation2 |
| CASSRGGXYEQYF | SSQRGGIYE | 4 | simulation1 |
| CASSRKGQGXNTEAFF | SRKGGGMWTE | 2 | simulation7 |
| CASSRNXGSTDTQYF | SRAAGSKDT | 2 | simulation6 |
| CASSRPGXGMNTEAFF | SRKGGGMWTE | 2 | simulation7 |
| CASSRXAGSGNTIYF | SRAAGSKDT | 2 | simulation6 |
| CASSRXAGXTDTQYF | SRAAGSKDT | 7 | simulation6 |
| CASSRXGGGXNTEAFF | SRKGGGMWTE | 4 | simulation7 |
| CASSRXGGMNTEAFF | SYERGMNTE | 2 | simulation2 |
| CASSRXGGRMNTEAFF | SRKGGGMWTE | 2 | simulation7 |
| CASSRXGGWYNEQFF | SRAGADYNE | 2 | simulation2 |
| CASSRXGGXMNTEAFF | SRKGGGMWTE | 4 | simulation7 |

| | | | |
|---|---|---|---|
| CASSRXGXTYNEQFF | SRAGADYNE | 3 | simulation2 |
| CASSRXLAGGAYEQYF | RTLLGGASE | 2 | simulation4 |
| CASSRXQGGMNTEAFF | SRKGGGMWTE | 4 | simulation7 |
| CASSRXXGSTDTQYF | SRAAGSKDT | 8 | simulation6 |
| CASSSEXGYGYTF | SYSEQGYG | 2 | simulation5 |
| CASSSLAGXHNEQFF | SHSLAGGSNE | 2 | simulation6 |
| CASSSLAGXSYEQYF | SHSLAGGSNE | 2 | simulation6 |
| CASSSLXTGLYNEQFF | KGTGLYNE | 2 | simulation2 |
| CASSSRAGGXNEQFF | SHSLAGGSNE | 2 | simulation6 |
| CASSSRLAGGXNEQFF | SHSLAGGSNE | 2 | simulation6 |
| CASSSRQGXYEQYF | SSQRGGIYE | 2 | simulation1 |
| CASSSRTSGSXDTQYF | SRAAGSKDT | 2 | simulation6 |
| CASSTGXRGTDTQYF | TGNRGADT | 3 | simulation5 |
| CASSTVXGGIYEQYF | SSQRGGIYE | 2 | simulation1 |
| CASSTXIGGSYEQYF | TPWGGSYE | 2 | simulation6 |
| CASSVLAGVXEQFF | VLAPVQE | 2 | simulation4 |
| CASSVVLAXVYEQYF | VLAPVQE | 2 | simulation4 |
| CASSWTGEKXYEQYF | WTGEKHE | 2 | simulation1 |
| CASSWTGXXYEQYF | SSQRGGIYE,WTGEKHE | 8 | simulation1 |
| CASSXAGSPDTQYF | SRAAGSKDT | 3 | simulation6 |
| CASSXAGTPYNEQFF | SRAGADYNE | 2 | simulation2 |
| CASSXAXGSTDTQYF | SRAAGSKDT | 7 | simulation6 |
| CASSXDPTGDXYEQYF | NPTGDFYE | 3 | simulation8 |
| CASSXEGDMNTEAFF | SLEGDMDSE | 2 | simulation8 |
| CASSXEQGYGYTF | SYSEQGYG | 5 | simulation5 |
| CASSXFAGANTEAFF | PFAGADT | 2 | simulation5 |
| CASSXGAPYNEQFF | SRAGADYNE | 2 | simulation2 |
| CASSXGAXYNEQFF | SRAGADYNE | 5 | simulation2 |
| CASSXGDRGADTQYF | TGNRGADT | 2 | simulation5 |
| CASSXGGGSYEQYF | SSQRGGIYE | 2 | simulation1 |
| CASSXGGIYEQYF | SSQRGGIYE | 2 | simulation1 |
| CASSXGGPPYNEQFF | PIPPYNE | 2 | simulation4 |
| CASSXGLAGADTQYF | PFAGADT | 3 | simulation5 |
| CASSXGLAGVPYEQYF | SPLLAGGPYE | 2 | simulation8 |
| CASSXGPGTGPYNEQFF | KGTGLYNE | 2 | simulation2 |
| CASSXGRGXYNEQFF | VVGRGAYNE | 10 | simulation7 |
| CASSXGSYEQYF | SRGSAET | 2 | simulation3 |
| CASSXGTGMNTEAFF | SYERGMNTE | 2 | simulation2 |
| CASSXGTGXYNEQFF | KGTGLYNE | 14 | simulation2 |
| CASSXGXDYNEQFF | SRAGADYNE | 3 | simulation2 |
| CASSXGXRGADTQYF | TGNRGADT | 6 | simulation5 |
| CASSXKGGGSYNEQFF | KGTGLYNE | 2 | simulation2 |

| | | | |
|---|---|---|---|
| CASSXKGXGSYNEQFF | KGTGLYNE | 3 | simulation2 |
| CASSXLAGGSNQPQHF | SHSLAGGSNE | 2 | simulation6 |
| CASSXLAGGYNEQFF | SHSLAGGSNE | 16 | simulation6 |
| CASSXLAGSTDTQYF | SRAAGSKDT | 10 | simulation6 |
| CASSXLAGXQETQYF | VLAPVQE | 11 | simulation4 |
| CASSXLGRAYGYTF | HLYRAYG | 2 | simulation1 |
| CASSXLQGINQPQHF | ELSGINQP | 2 | simulation3 |
| CASSXLRGMNTEAFF | SYERGMNTE | 2 | simulation2 |
| CASSXLSGRNQPQHF | ELSGINQP | 4 | simulation3 |
| CASSXLSGXNQPQHF | ELSGINQP | 3 | simulation3 |
| CASSXLTGINQPQHF | ELSGINQP | 4 | simulation3 |
| CASSXLYRGYGYTF | HLYRAYG | 2 | simulation1 |
| CASSXNPTGGSYEQYF | NPTGDFYE | 2 | simulation8 |
| CASSXPGGHYEQYF | SSQRGGIYE | 2 | simulation1 |
| CASSXPGLAGADTQYF | PFAGADT | 2 | simulation5 |
| CASSXPGLAGGPYEQYF | SPLLAGGPYE | 2 | simulation8 |
| CASSXPGLPYNEQFF | PIPPYNE | 2 | simulation4 |
| CASSXPLAGGPYNEQFF | SPLLAGGPYE | 2 | simulation8 |
| CASSXPPGPYNEQFF | PIPPYNE | 2 | simulation4 |
| CASSXPTGDXYEQYF | NPTGDFYE | 9 | simulation8 |
| CASSXPTGGTDTQYF | SEPTSGRDT | 2 | simulation7 |
| CASSXPTGPYNEQFF | PIPPYNE | 2 | simulation4 |
| CASSXPTGXFYEQYF | NPTGDFYE | 7 | simulation8 |
| CASSXPXGGSYEQYF | TPWGGSYE | 6 | simulation6 |
| CASSXQLAGVQETQYF | VLAPVQE | 2 | simulation4 |
| CASSXRDRGAYNEQFF | VVGRGAYNE | 2 | simulation7 |
| CASSXREQGYGYTF | SYSEQGYG | 2 | simulation5 |
| CASSXRGAYNEQFF | VVGRGAYNE | 4 | simulation7 |
| CASSXRGGAPYNEQFF | SRAGADYNE | 2 | simulation2 |
| CASSXRGSSYEQYF | SSQRGGIYE | 2 | simulation1 |
| CASSXRGSYEQYF | SRGSAET | 2 | simulation3 |
| CASSXRGXAYEQYF | SSQRGGIYE | 3 | simulation1 |
| CASSXRQGSYEQYF | SSQRGGIYE | 4 | simulation1 |
| CASSXRTGLYNEQFF | KGTGLYNE | 3 | simulation2 |
| CASSXRTGSYEQYF | SSQRGGIYE | 2 | simulation1 |
| CASSXRWTGEKLFF | WTGEKHE | 2 | simulation1 |
| CASSXSGGXYEQYF | SSQRGGIYE | 6 | simulation1 |
| CASSXSGINQPQHF | ELSGINQP | 2 | simulation3 |
| CASSXSRLAGITDTQYF | SRAAGSKDT | 2 | simulation6 |
| CASSXSXXGYGYTF | SYSEQGYG | 57 | simulation5 |
| CASSXTGDFYEQYF | NPTGDFYE | 5 | simulation8 |
| CASSXTGLYNEQFF | KGTGLYNE | 3 | simulation2 |

| | | | |
|---|---|---|---|
| CASSXTGTGGSYEQYF | TPWGGSYE | 2 | simulation6 |
| CASSXTGTRGTDTQYF | TGNRGADT | 4 | simulation5 |
| CASSXTLAGGAYEQYF | RTLLGGASE | 2 | simulation4 |
| CASSXTSGRDNEQFF | SEPTSGRDT | 2 | simulation7 |
| CASSXTSGRDSTDTQYF | SEPTSGRDT | 2 | simulation7 |
| CASSXTSGXETQYF | SEPTSGRDT | 6 | simulation7 |
| CASSXTXGGGSYEQYF | TPWGGSYE | 3 | simulation6 |
| CASSXVGGYEQYF | SLAVGGYE | 3 | simulation3 |
| CASSXVLAGXQETQYF | VLAPVQE | 5 | simulation4 |
| CASSXVLAXGQETQYF | VLAPVQE | 3 | simulation4 |
| CASSXWTGEKLFF | WTGEKHE | 7 | simulation1 |
| CASSXXAGXDTQYF | PFAGADT | 24 | simulation5 |
| CASSXXGDRGADTQYF | TGNRGADT | 4 | simulation5 |
| CASSXXGGGMNTEAFF | SRKGGGMWTE | 27, 3 | simulation7 |
| CASSXXGGXYEQYF | SSQRGGIYE | 10 | simulation1 |
| CASSXXGTGXYNEQFF | KGTGLYNE | 5 | simulation2 |
| CASSXXGXNTEAFF | SYERGMNTE | 44 | simulation2 |
| CASSXXLAGGSNEQFF | SHSLAGGSNE | 3 | simulation6 |
| CASSXXLAGGSYEQYF | SPLLAGGPYE,SHSLAGGSNE | 8, 6 | simulation6,simulation8 |
| CASSXXLAGGYNEQFF | SHSLAGGSNE | 8 | simulation6 |
| CASSXXLAGVQETQYF | VLAPVQE | 8 | simulation4 |
| CASSXXTSGTDTQYF | SEPTSGRDT | 3 | simulation7 |
| CASSXYRAYGYTF | HLYRAYG | 6 | simulation1 |
| CASSYAGAXYNEQFF | SRAGADYNE | 2 | simulation2 |
| CASSYLAGXPYEQYF | SPLLAGGPYE | 2 | simulation8 |
| CASSYSEGXYGYTF | SYSEQGYG | 2 | simulation5 |
| CASSYSEQGXYGYTF | SYSEQGYG | 2 | simulation5 |
| CASSYSRQGXGYGYTF | SYSEQGYG | 2 | simulation5 |
| CASSYSXGLNTEAFF | SYERGMNTE | 2 | simulation2 |
| CASSYSXQGYEQYF | SYSEQGYG | 4 | simulation5 |
| CASSYSXQVYGYTF | SYSEQGYG | 2 | simulation5 |
| CASSYSXQXYGYTF | SYSEQGYG | 3 | simulation5 |
| CASSYXGLAGVQETQYF | VLAPVQE | 2 | simulation4 |
| CASSYXKQGYGYTF | SYSEQGYG | 2 | simulation5 |
| CASSYXTGGNTEAFF | SYERGMNTE | 3 | simulation2 |
| CASTLXGGAYEQYF | RTLLGGASE | 3 | simulation4 |
| CASTPGXGSYEQYF | TPWGGSYE | 4 | simulation6 |
| CASTPLGXSYEQYF | TPWGGSYE | 2 | simulation6 |
| CASTPXGSSYEQYF | TPWGGSYE | 2 | simulation6 |
| CASTPXXGSYEQYF | TPWGGSYE | 7 | simulation6 |
| CASTVWGXSYEQYF | TPWGGSYE | 2 | simulation6 |
| CASTXRGGSYEQYF | TPWGGSYE | 3 | simulation6 |

| | | | |
|---|---|---|---|
| CASTXTGGSYEQYF | TPWGGSYE | 2 | simulation6 |
| CASTXWGSSYEQYF | TPWGGSYE | 2 | simulation6 |
| CASXASGGYEQYF | SLAVGGYE | 2 | simulation3 |
| CASXAXGGYEQYF | SLAVGGYE | 7 | simulation3 |
| CASXEGSMNTEAFF | SYERGMNTE | 2 | simulation2 |
| CASXGGGXYNEQFF | KGTGLYNE,SRAGADYNE | 7 | simulation2 |
| CASXGLQETQYF | SRGSAET | 2 | simulation3 |
| CASXGRGPYNEQFF | VVGRGAYNE | 2 | simulation7 |
| CASXGTGXYNEQFF | KGTGLYNE | 8 | simulation2 |
| CASXGXGLYNEQFF | KGTGLYNE | 3 | simulation2 |
| CASXGXRGADTQYF | TGNRGADT | 6 | simulation5 |
| CASXKLGGPPYNEQFF | PIPPYNE | 2 | simulation4 |
| CASXLAGGPYEQYF | SPLLAGGPYE | 2 | simulation8 |
| CASXLAXXQETQYF | VLAPVQE | 18 | simulation4 |
| CASXLTSGXDTQYF | SEPTSGRDT | 3 | simulation7 |
| CASXLXGGAXEQFF | RTLLGGASE | 6 | simulation4 |
| CASXLXRAYGYTF | HLYRAYG | 15 | simulation1 |
| CASXPFEGTDTQYF | PFAGADT | 2 | simulation5 |
| CASXPGWDYNEQFF | SRAGADYNE | 2 | simulation2 |
| CASXPTGDXYEQYF | NPTGDFYE | 13 | simulation8 |
| CASXPTGXFYEQYF | NPTGDFYE | 6 | simulation8 |
| CASXPTSGRCTDTQYF | SEPTSGRDT | 2 | simulation7 |
| CASXRLAGGSGELFF | SHSLAGGSNE | 2 | simulation6 |
| CASXRLAGSTDTQYF | SRAAGSKDT | 2 | simulation6 |
| CASXRVGGYEQYF | SLAVGGYE | 2 | simulation3 |
| CASXSGINQPQHF | ELSGINQP | 3 | simulation3 |
| CASXSXQGYGYTF | SYSEQGYG | 10 | simulation5 |
| CASXTLAGGAYEQYF | RTLLGGASE | 2 | simulation4 |
| CASXWTGEKLFF | WTGEKHE | 2 | simulation1 |
| CASXWTGENTEAFF | WTGEKHE | 4 | simulation1 |
| CASXXAGADTQYF | PFAGADT | 8 | simulation5 |
| CASXXAGSTDTQYF | SRAAGSKDT | 12 | simulation6 |
| CASXXGGGMNTEAFF | SRKGGGMWTE | 22 | simulation7 |
| CASXXPPYNEQFF | PIPPYNE | 3 | simulation4 |
| CASXYEGSGSNQPQHF | ELSGINQP | 2 | simulation3 |
| CASXYRAYGYTF | HLYRAYG | 2 | simulation1 |
| CATGTXGADTQYF | TGNRGADT | 2 | simulation5 |
| CATGXPGADTQYF | TGNRGADT | 2 | simulation5 |
| CATLYRXYGYTF | HLYRAYG | 2 | simulation1 |
| CATPXGGAYEQYF | TPWGGSYE | 2 | simulation6 |
| CATPXGSSYEQYF | TPWGGSYE | 2 | simulation6 |
| CATRXRGADTQYF | TGNRGADT | 2 | simulation5 |

| | | | |
|---|---|---|---|
| CATSDXGADTQYF | PFAGADT,TGNRGADT | 2 | simulation5 |
| CATSRAEGLXDTQYF | SRAAGSKDT | 2 | simulation6 |
| CATSRXGGGXNTEAFF | SRKGGGMWTE | 3 | simulation7 |
| CATSRXGRGMNTEAFF | SRKGGGMWTE | 2 | simulation7 |
| CAWRDXLAGGPYEQYF | SPLLAGGPYE | 2 | simulation8 |
| CAWTGEXYEQYF | WTGEKHE | 2 | simulation1 |
| CAWXPGGPYNEQFF | PIPPYNE | 2 | simulation4 |
| CAXGLAGGGYEQYF | SLAVGGYE | 2 | simulation3 |
| CAXLAGVQETQYF | VLAPVQE | 3 | simulation4 |
| CAXSELGGXNQPQHF | ELSGINQP | 3 | simulation3 |
| CAXSEQGYGYTF | SYSEQGYG | 2 | simulation5 |
| CAXSGINQPQHF | ELSGINQP | 2 | simulation3 |
| CAXSPXGGADTQYF | PFAGADT | 3 | simulation5 |
| CAXSRRGGSTDTQYF | SRAAGSKDT | 2 | simulation6 |
| CAXSTGEKKETQYF | WTGEKHE | 2 | simulation1 |
| CAXWTGEKLFF | WTGEKHE | 2 | simulation1 |
| CAXWTGEXYEQYF | WTGEKHE | 3 | simulation1 |
| CSAPTGXFYEQYF | NPTGDFYE | 2 | simulation8 |
| CSARGXDYNEQFF | SRAGADYNE | 2 | simulation2 |
| CSARLSGXNQPQHF | ELSGINQP | 2 | simulation3 |
| CSARXLAGGAYEQYF | RTLLGGASE | 2 | simulation4 |
| CSASXQGYGYTF | SYSEQGYG | 2 | simulation5 |
| CSAXLDRAYGYTF | HLYRAYG | 2 | simulation1 |
| CSAXRGRNTEAFF | SYERGMNTE | 3 | simulation2 |
| CSAXYRAYGYTF | HLYRAYG | 2 | simulation1 |
| CSXARGGYEQYF | SLAVGGYE | 2 | simulation3 |
| CXWTGEKLFF | WTGEKHE | 2 | simulation1 |

**Table S6.3:** Overview of the CDR3 β content of the clusters highlighted by yellow rows in table S6.2.

| Consensus motif | LIgO seed | Sequences | Simulation |
|---|---|---|---|
| CASSPGLAGGXNEQFF | SHSLAGGSNE | CASSPGLAGGDNEQFF, CASSPGLAGGQNEQFF, CASSPGLAGGFNEQFF, CASSPGLAGGINEQFF | Simulation 6 |
| CASSPGLAGGXNEQFF | SPLLAGGPYE | CASSPGLAGGDNEQFF, CASSPGLAGGQNEQFF, CASSPGLAGGINEQFF | Simulation 8 |
| CASSXXLAGGSYEQYF | SHSLAGGSNE | CASSQGLAGGSYEQYF, CASSYGLAGGSYEQYF, CASSRGLAGGSYEQYF, CASSRRLAGGSYEQYF, CASSRLLAGGSYEQYF, CASSQELAGGSYEQYF, CASSLRLAGGSYEQYF, CASSLVLAGGSYEQYF | Simulation 6 |
| CASSXXLAGGSYEQYF | SPLLAGGPYE | CASSRGLAGGSYEQYF, CASSRRLAGGSYEQYF, CASSQELAGGSYEQYF, CASSQGLAGGSYEQYF, CASSLRLAGGSYEQYF, CASSLVLAGGSYEQYF | Simulation 8 |

**S7: UMAP of LIgO simulated repertoires**

Figure 4B in the main text contains a UMAP plot of one simulation. Following seven figures contain the UMAP plots of the additional seven TCR repertoires that were simulated by LIgO. The CDR3 β sequences that were shared between different simulations are not shown in these individual UMAPs.

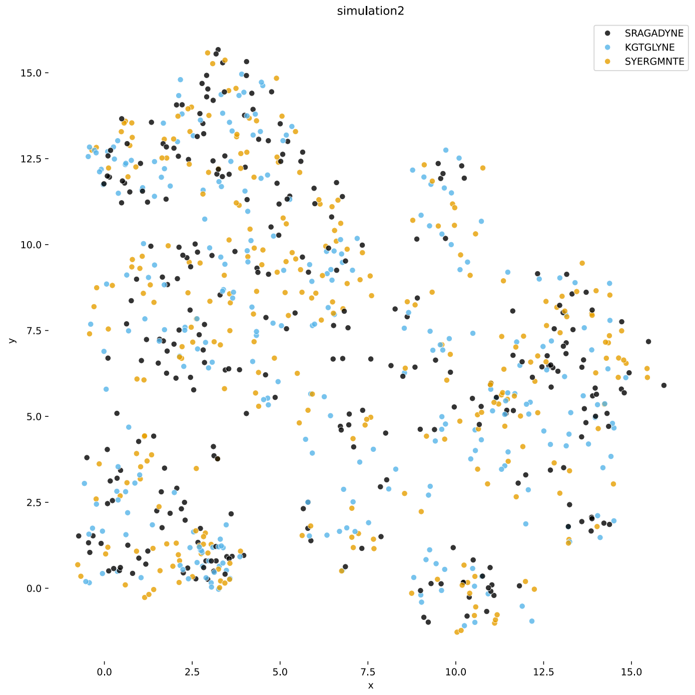


**Figure S7.1:** UMAP plot of simulation 2, colored by LIgO seed.

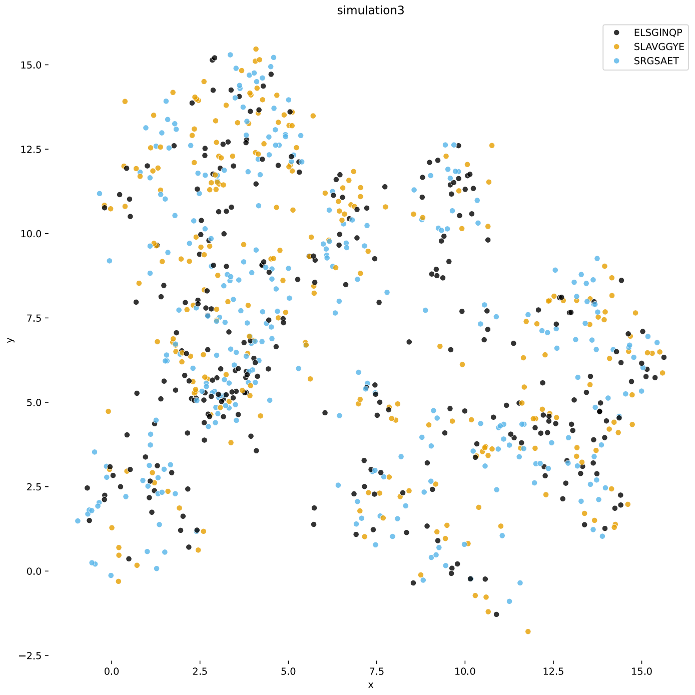


**Figure S7.2:** UMAP plot of simulation 3, colored by LIgO seed.

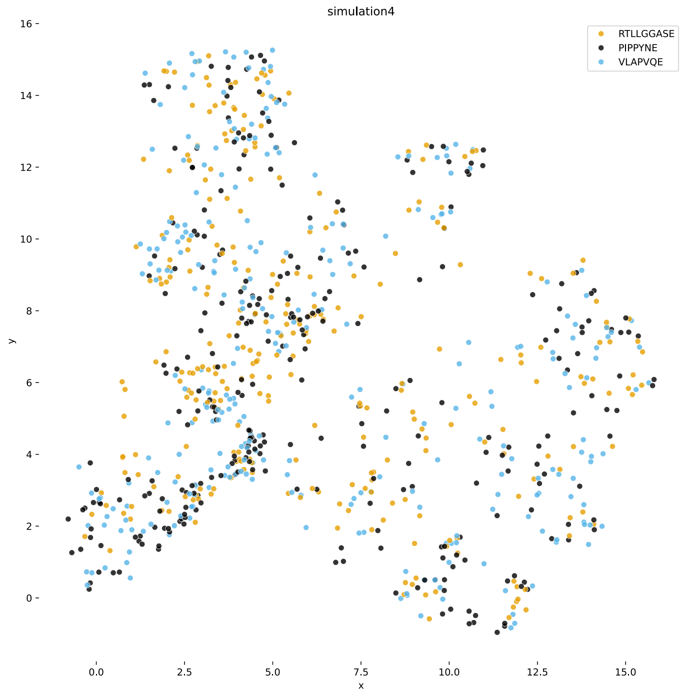


**Figure S7.3:** UMAP plot of simulation 4, colored by LIgO seed.

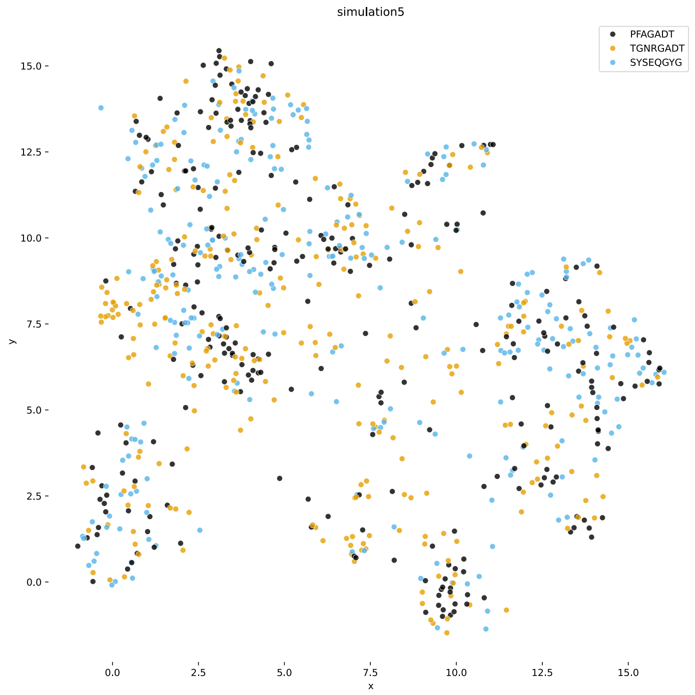


**Figure S7.4:** UMAP plot of simulation 5, colored by LIgO seed.

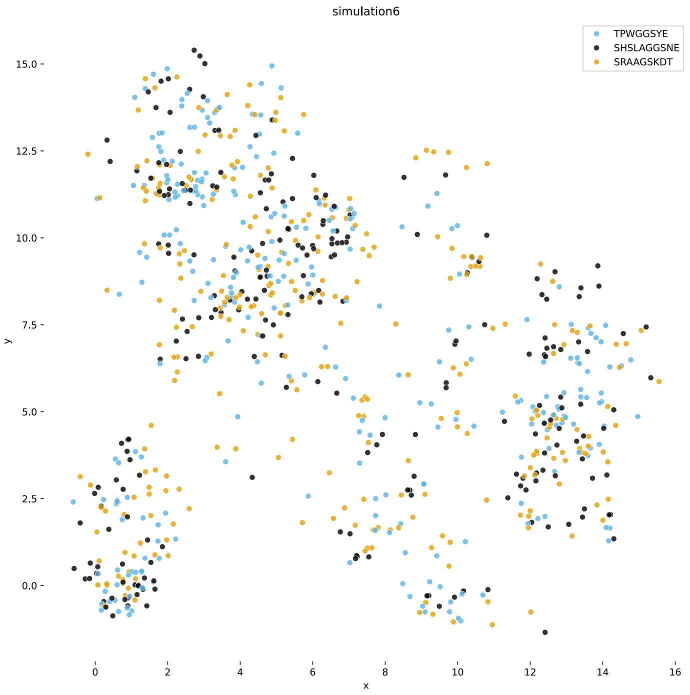


**Figure S7.5:** UMAP plot of simulation 6, colored by LIgO seed.

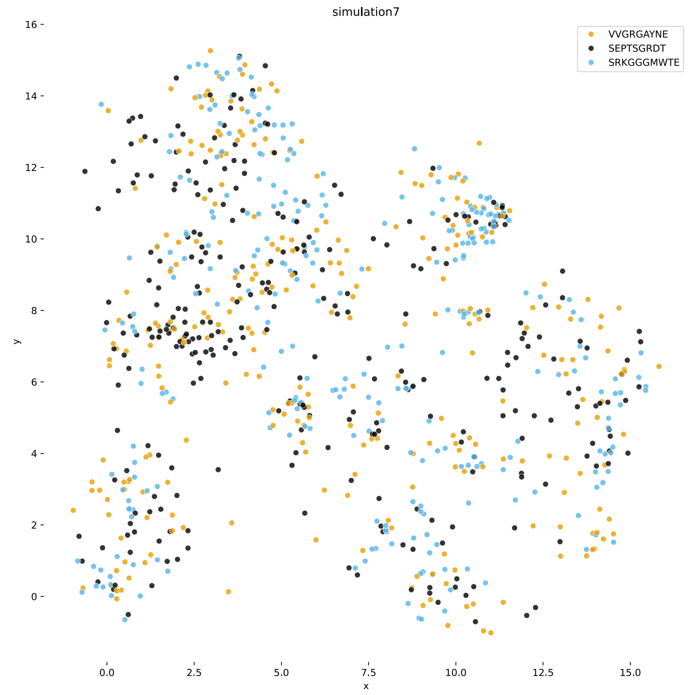


**Figure S7.6:** UMAP plot of simulation 7, colored by LIgO seed.

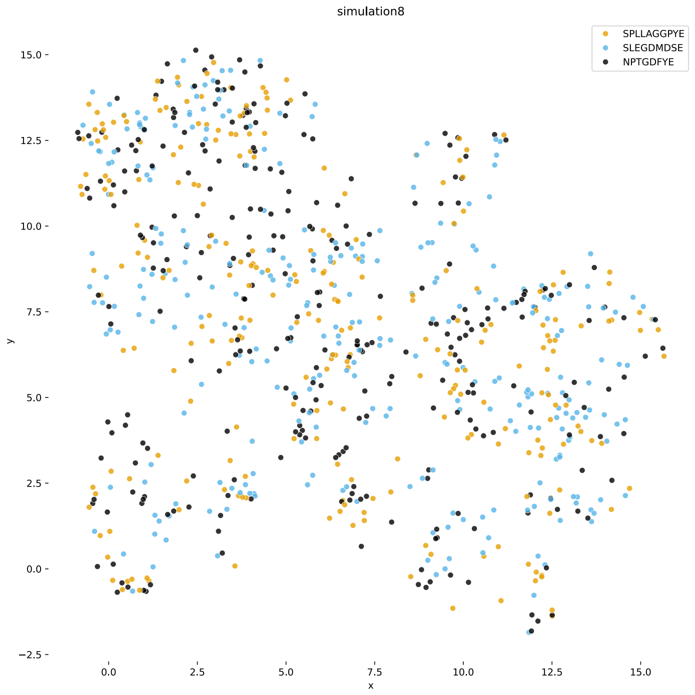


**Figure S7.7:** UMAP plot of simulation 8, colored by seed.

**S8: Overview of the generation probability for clustered TCRs, singlets and new linking TCRs**

Figure 5 in the main text gives an overview of the generation probability for the clustered TCRs, singlets and linking TCRs for one experimental and simulated TCR dataset. Figures S8.1-S8.4 list the results for all studied experimental and simulated TCR repertoires.

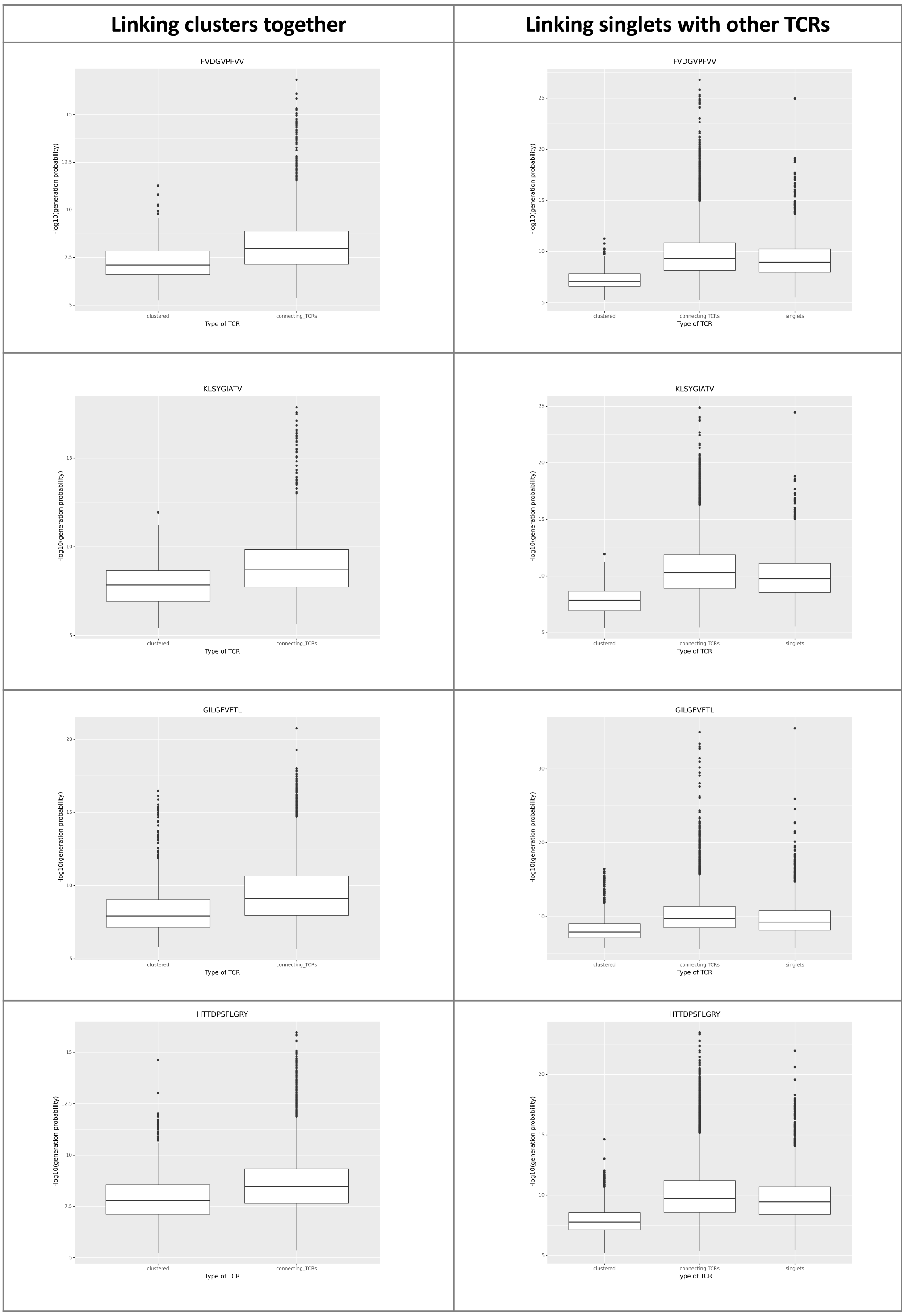


**Figure S8.1:** Overview of the generation probability for clustered TCRs, singlets and new linking TCRs for four studied experimental epitope-specific TCR repertoires.

| Linking clusters together | Linking singlets with other TCRs |
| --- | --- |

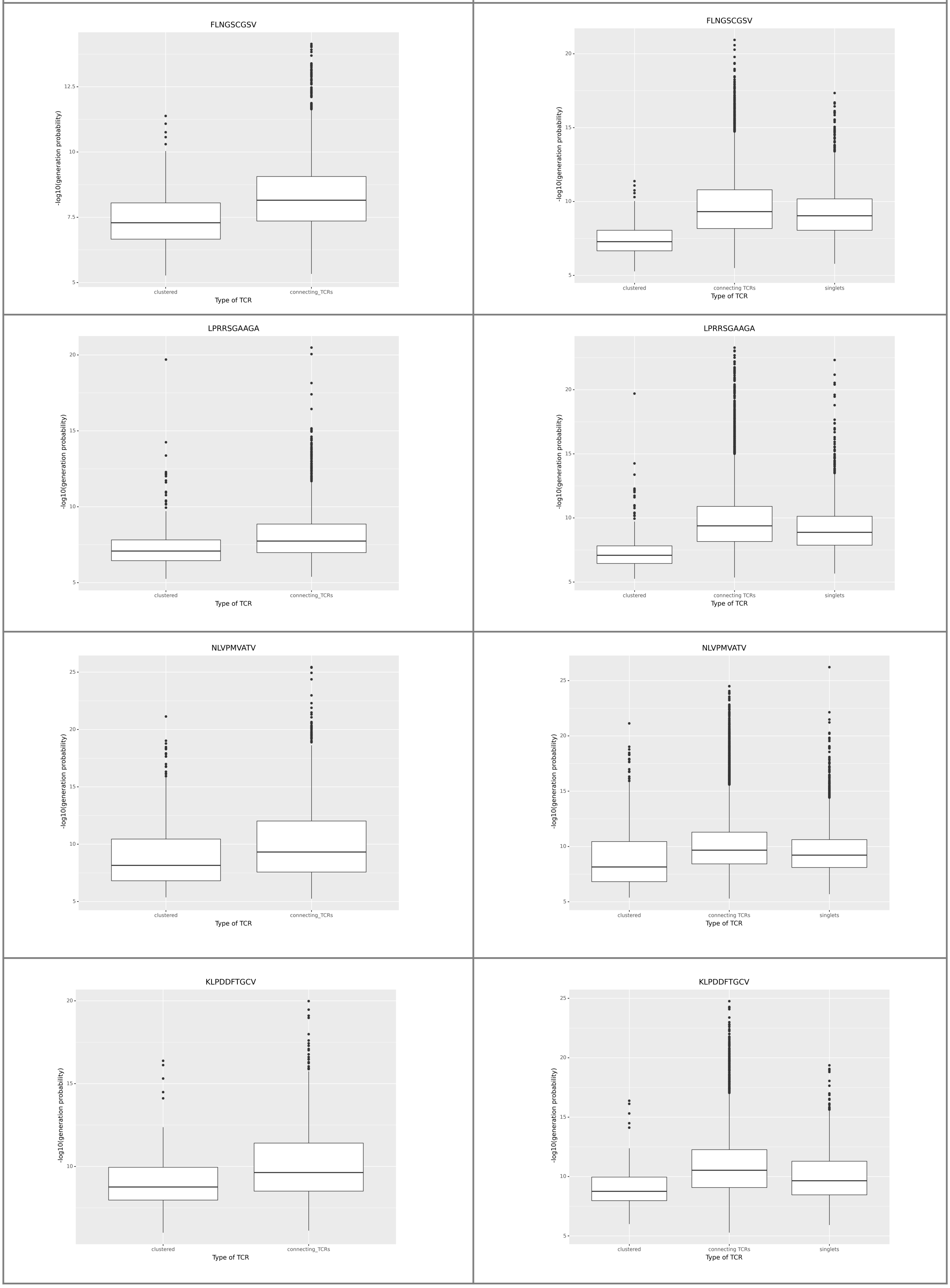


**Figure S8.2:** Overview of the generation probability for clustered TCRs, singlets and new linking TCRs for the four remaining studied experimental epitope-specific TCR repertoires

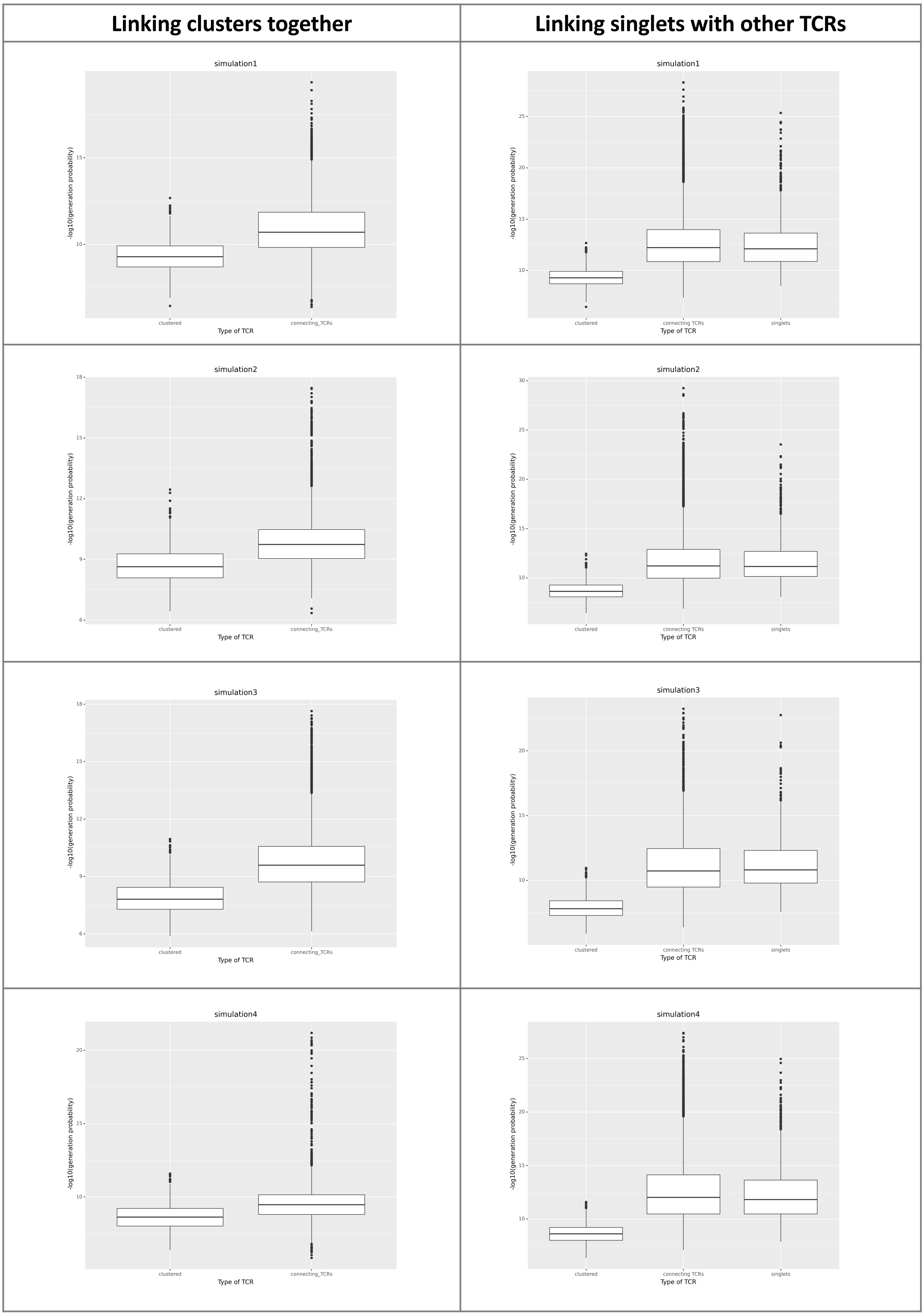


**Figure S8.3:** Overview of the generation probability for clustered TCRs, singlets and new linking TCRs for the first four studied simulated TCR repertoires.

| Linking clusters together | Linking singlets with other TCRs |
|---|---|

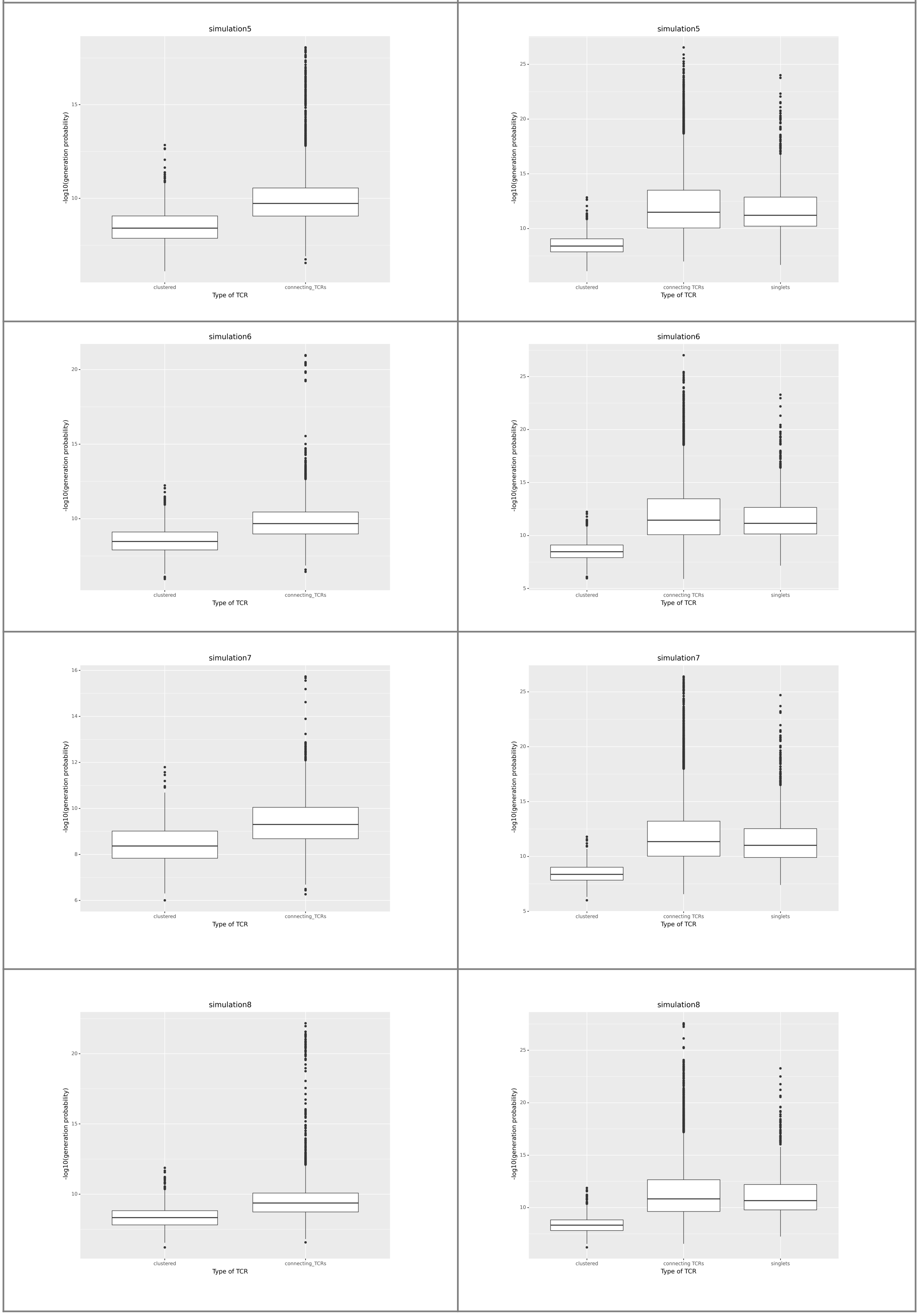


**Figure S8.4:** Overview of the generation probability for clustered TCRs, singlets and new linking TCRs for the remaining four studied simulated TCR repertoires.
